# No evidence for direct physical interaction of 5-HT_2A_-mGluR2 receptors *in vitro* or *in vivo*

**DOI:** 10.64898/2026.06.28.734515

**Authors:** Blake A. Fordyce, Yi-Ting Chiu, Nicholas J. Wright, Kensuke Sakamoto, Scott P. Lyons, Thomas S. Webb, Hayleigh E. Tilton, Jessica J. Walsh, Gerard Marek, Vincent Setola, Bryan L. Roth

**Author notes:** Gilgamesh Pharma Inc., New York, NY 10003, USA.

## Abstract

Activation of mGluR2 (metabotropic glutamate receptor 2), the primary presynaptic autoreceptor for glutamate in the brain, is well established to attenuate the psychedelics-mediated behavioral and electrophysiological effects. However, the mechanisms responsible for these actions are controversial. The two competing mechanistic hypotheses have been proposed to explain this phenomenon are: (1) direct actions mediated by mGluR2/5-HT_2A_ heterodimers, and (2) inhibition of 5-HT_2A_-mediated excitation of pyramidal neurons via presynaptic inhibition of glutamate release by mGluR2 receptors. Consistent with prior reports, we show that mGluR2 agonist pretreatment attenuates the head twitch response induced by the psychedelic drug 1-(2,5-Dimethoxy-4-iodophenyl)-2-aminopropane (DOI) in mice engineered to express mGluR2-mCherry and 5-HT_2A_-eGFP-CT tagged receptors. We next employed multiple orthogonal *in vivo* and *in vitro* approaches to explore the potential for direct physical interactions between mGluR2 and 5-HT_2A_ receptors. Across all approaches, we found no evidence for receptor colocalization or oligomerization under basal or 5-HT_2A_ agonist-exposed conditions *in vitro* or *in vivo*. Radioligand binding and kinetic analyses revealed no evidence for mGluR2-mediated modulation of 5-HT_2A_ ligand binding *in vitro* or *in vivo*. Collectively, our findings support models in which mGluR2 signaling modulates the activity of Gα_q_-coupled 5-HT_2A_ receptors in layer V pyramidal neurons, rather than models positing the requirement of mGluR2/5-HT_2A_ multimers.

## Introduction

Current pharmacological treatments for neuropsychiatric disorders, including those for schizophrenia, depression, and post-traumatic stress disorder, frequently target serotonin (5-hydroxytryptamine; 5-HT) receptors for their actions^1^. Among the various 5-HT receptors, the 5-HT_2A_ serotonin receptor (5-HT_2A_) receptor represents a major target for neuropsychiatric drugs including psychedelic, antidepressant and antipsychotic drugs^1–4^. Despite their complex polypharmacology^5^, psychedelic actions are mediated by 5-HT_2A_ receptor activation^6–8^, as demonstrated by their abolition in 5-HT_2A_ knockout mice and blockade by 5-HT_2A_ antagonists. Furthermore, multiple randomized, placebo-controlled Phase 2 and 3 clinical trials have demonstrated that one or two administrations of psilocybin produce sustained and statistically-significant reductions in depressive symptoms^9–11^.

Psychedelics are well-known to enhance glutamatergic neurotransmission in the cortex, a phenomenon first demonstrated by whole-cell patch-clamp recordings in 1997^12^. These pioneering studies disclosed an enhancement of excitatory postsynaptic potentials (EPSPs) in layer V pyramidal neurons of the medial prefrontal cortex (mPFC) upon psychedelic drug administration *ex vivo*^12^. More recent studies in mice have established that these effects are mediated by 5-HT_2A_ receptor activation of Gα_q_ signaling^13^. Importantly, orthosteric mGluR2/3 antagonists (e.g., LY341495) amplify the frequency and amplitude of 5-HT_2A_-mediated excitatory postsynaptic currents (EPSCs), which are, conversely, suppressed by mGluR2/3 agonists. As a group II metabotropic glutamate receptor member of the class C G-Protein Coupled Receptor (GPCR) family, mGluR2 couples to G_i/o/z_-type Gα subunits^14^ to inhibit presynaptic glutamate release. mGluRs feature a large extracellular Venus flytrap (VFT) domain at the N-terminus that contains the orthosteric binding site and is key for their obligate dimerization^15,16^.

These prior electrophysiology results are consistent with mechanisms whereby mGluR2/3 receptors modulate cortical excitability of layer V neurons either directly^17–19^ or indirectly^20–22^. In rodents and non-human primates, mGluR2 is predominantly expressed presynaptically in glutamatergic neurons, whereas mGluR3 shows broader distribution including neuronal and astrocytic compartments^23,24^. 5-HT_2A_ receptors, on the other hand, are localized to postsynaptic domains on Layer V cortical glutamatergic neurons^25–28^.

Circuit-level studies reveal a complex interplay between 5-HT_2A_ and mGluR2 receptors. Lesions to the medial thalamus, a major glutamatergic afferent to the mPFC, reduce 5-HT–evoked EPSCs, decrease mGluR2/3 receptor density, and paradoxically increase 5-HT_2A_ receptor density within the mPFC^29^. These results can be explained by a loss of thalamocortical glutamatergic drive to the mPFC, resulting in compensatory upregulation of postsynaptic 5-HT_2A_ receptors, while decreased mGluR2/3 receptor density reflects a direct loss of thalamocortical afferents. This explanation aligns with subsequent anatomical findings, demonstrating 5-HT_2A_ expression at postsynaptic sites within the cortex^26^.

Consistent with these findings, mGluR2/3 agonists suppress DOI-induced head-twitch responses (HTR) in rodents, while antagonists enhance the HTR^21,30,31^. Notably, mGluR2 knockout mice exhibit abolished DOI-induced HTR, highlighting a specific role for mGluR2 in modulating psychedelic-induced behaviors^32^. This counter-intuitive result points to a strong functional convergence of the 5-HT_2A_ and mGluR2 receptors, required for full hallucinogenic-like behaviors. Together, these findings have been interpreted to support a model in which presynaptic mGluR2 autoreceptors regulate postsynaptic 5-HT_2A_ receptor activity in layer V pyramidal neurons by inhibition of glutamate release^18,32–35^.

A competing model posits that 5-HT_2A_ receptors engage transmembrane domain 4 (TM4) of mGluR2 receptors to assemble functional heterodimers in cortical neurons, thereby potentiating G_i/o/z_-coupled signaling in response to psychedelic compounds^20,21^. Although compelling, this model contrasts with the known and well-established canonical subcellular localization and physiological actions of 5-HT_2A_ and mGluR2^13,15,16,23–28^. Directly examining the distribution and physical association between 5-HT_2A_ and mGluR2 receptors represents a first step towards resolving these apparent inconsistencies.

In addition to models based on physical association, prior studies have also suggested that activation of mGluR2/3 receptors may alter agonist binding affinity at 5-HT_2A_ receptors in native tissue^21^, raising the possibility of receptor-level allosteric modulation independent of stable heterodimer formation. This interpretation predicts that mGluR2 activation should measurably shift 5-HT_2A_ ligand binding properties or alter ligand binding kinetics. Directly testing these possibilities provides a complementary approach to distinguish receptor-level mechanisms from models in which mGluR2 signaling modulates 5-HT_2A_ function indirectly at the circuit level.

To test these competing hypotheses, we created mice expressing fluorescently tagged 5-HT_2A_ (EGFP) (Chiu et al., *Nature Neurosci in press*) and mGluR2 (mCherry) receptors^14^. Using a variety of orthogonal *in vivo* and *in vitro* approaches, we confirm that 5-HT_2A_ receptors are localized to the apical dendrites of layer V pyramidal neurons, while mGluR2 receptors are largely found in the neuropil and layers IV/V of the cortex, consistent with many previous findings^23,25,26,36,37^. Significantly, we find no evidence of 5-HT_2A_–mGluR2 colocalization or functional heterodimerization under basal or 5-HT_2A_ agonist-exposed conditions. Instead, these results support classical models of prefrontal cortical microcircuitry in which presynaptic mGluR2 receptors *indirectly* modulate postsynaptic 5-HT_2A_ receptor activity in layer V neurons.

## Methods and materials

### Animals

Adult Htr2a-EGFP-CT-IRES-CreERT2 (*Htr2a*^EGFP-CreERT2^; Htr2a-EGFP-CT)^38^ and mGluR2-mCherry-CT-Flpo (Grm2^mCherry-Flpo^, mGluR2-mCherry)^14^ mice were cross bred in house to create Htr2a-EGFP-CT X mGluR2-mCherry offspring which were used in this study at 8-20 weeks of age.

Separate lines of transgenic mice with a deletion of mGluR2 (Taconic Farms, Germantown, NY) were generated by homologous recombination^39^. *Htr2a* knockout (*Htr2a*^−/−^) mice, and their respective wild-type littermates were bred from heterozygous breeders obtained as previously described^40^.

Mice were housed 3–5 mice per cage according to sex and genotype in a light-dark controlled room under a 12hr–12hr cycle (lights on at 07:00) with food and water provided ad libitum. All experiments were conducted during the light cycle with an approved protocol from the University of North Carolina at Chapel Hill Institutional Animal Care and Use Committee and were performed in accordance with the National Institutes of Health Guide for Care and Use of Laboratory Animals and as approved by the Animal Care and Use Committee of the University of North Carolina (UNC).

### Drug administration

The drugs used in the experiments were serotonin (Tocris Bioscience, Bristol, UK), lysergic acid diethylamide (LSD; Sigma-Aldrich, St. Louis, MO, USA), 25CN-NBOH (Axon Medchem, Groningen, The Netherlands), DOI (Bio-Techne Corp., Minneapolis, MN), LY379268 (Cayman Chemical, Ann Arbor, MI, USA), and LY341495 (Cayman Chemical, Ann Arbor, MI, USA). Stock solutions for pharmacology assays were prepared in sterile dH_2_0 whereas solutions for behavioral paradigms were prepared fresh in 0.9% saline. Compounds were administered to animals intraperitoneally at doses of 1 mg/kg for DOI and 15 mg/kg for LY379268, delivered in a volume of 10 mL/kg from stock solutions of 0.1 mg/mL and 1.5 mg/mL, respectively. Total volume injected was based on animal weight during the day of treatment.

### Head twitch response assay

Head twitch responses (HTRs) were measured using an automated magnetometer-based assay adapted from De la Fuente Revenga et. al^41^. Male and female Htr2a-EGFP-CT X mGluR2-mCherry mice (n=16 per group, age 10-20 weeks) were fitted with magnetic ear tags (La Pias Ear Tags, Stoelting Co.; N50 neodymium magnets, 3mm x 1mm) under brief isoflurane anesthesia one week prior to behavioral testing.

On testing days, mice were acclimated to the behavioral room for 1 hr before the start of the experiment. Animals were randomly assigned to treatment groups and experimenter was blinded to condition. Baseline HTR activity was measured following intraperitoneal administration (i.p.) of saline. For drug testing, mice received either saline or LY379268 (15 mg/kg, i.p.) 30 min prior to administration of DOI (1 mg/kg, i.p). HTRs were recorded for 15 min immediately following DOI administration.

Magnetometer signals were acquired using a NI USB-6001 data acquisition device (National Instruments) and analyzed using custom MATLAB (Mathworks, R2025a) scripts. To validate automated HTR detection, simultaneous video recordings were collected at 60 frames/sec and a subset of sessions was manually scored by an experimenter blinded to treatment condition.

### Tissue collection

Animals were intracardially perfused with ice cold 1X PBS and 4% PFA. Brain tissue was collected from animals at postnatal day 30-60, placed in 4% PFA overnight at 4°C after perfusion and transferred to 30% sucrose for the next 3 days. Tissue was sliced at 40 microns using a Lecia microtome and stored in anti-freeze media at -20°C.

### Primary cortical neurons

*Htr2a*^EGFP-CreERT2/EGFP-CreERT2^ and *Grm2*^mCherry-FlpO/mCherry-FlpO^ mice were crossed, and pregnant female mice were sacrificed at E16/17 to isolate cortical tissue from embryonic pups. Tissue was dissociated using papain (ThermoFisher Scientific, Waltham, MA, USA) and DNase (Sigma-Aldrich, St. Louis, MO, USA), incubated at 37°C for exactly 30 min. Papain solution was then neutralized with Glidia media (Dulbecco’s Modified Eagle Medium (DMEM; Corning, Corning, NY, USA), Fetal Bovine Serum (FBS; Genesee Scientific, San Diego, CA, USA), GlutaMAX (ThermoFisher Scientific, Waltham, MA, USA), Antibiotic-Antimycotic (ThermoFisher Scientific, Waltham, MA, USA,), spun down for 2 min at 4000 RPM, and washed with 1X HBSS. Neurons were then resuspended in Glidia media, filtered through a 70 *μ*Μ filter and plated in non-coated 6-well plates and placed in an incubator at 37 °C under 5% CO2 for 15 min to remove glial cells. Cells were then collected, counted and plated in seeding media (Neurobasal plus medium, Glutamax, Antibiotic-Antimycotic, B27+, (ThermoFisher Scientific, Waltham, MA, USA)) at 1.5 x 10^6^ cells/well in a poly-D-lysine coated 6-well or 0.5 x 10^6^ neurons per well using a 12-well plate with poly-D-lysine coated coverslips (Neuvitro Corporation, Vancouver, WA, USA). The following day, half of the seeding media was replaced with feeding media (Neurobasal plus medium, FDU, B27+, Glutamax). This feeding regimen was repeated every 2-3 days.

### RT-qPCR analysis of *Htr2a* and *Grm2 transcripts*

Total RNA was isolated from primary cortical neurons (B6 wild-type and double knock-in) at DIV 0, 2, 5, 10, 14, and 21. RNA concentrations and quality were measured using nanodrop and normalized across samples. One-step RT-qPCR was performed using TaqMan Fast Virus 1-Step Master Mix (ThermoFisher Scientific, Waltham, MA, USA) with FAM-labeled target probes (*Htr2a* or *Grm2*) and a VIC-labeled GAPDH probe. Each 20 µL reaction contained 50 ng RNA and was run in triplicate alongside no-template and no-reverse-transcription controls. Reactions were run on a Bio-Rad CFX96 (Bio-Rad Laboratories, Hercules, CA, USA) with standard cycling conditions. Relative expression was calculated using the ΔΔCq method normalized to DIV0 of each genotype, and fold changes were expressed as 2^(-ΔΔCq). Data are presented as mean ± SEM and analyzed using two-way ANOVA.

### Immunofluorescence microscopy

Primary cortical neurons were used for immunofluorescent microscopy at DIV 14,18, 21 and 28. Media was aspirated from the well, and washed with 1X PBS 3 times for 5 min. Neurons are then exposed to 4% PFA for 10 min at room temperature. The protocols for both primary cortical neuron cultures and whole brain tissue slices are identical. Tissue slices/cells are washed with 0.1% Triton-100 in 1X PBS 3 times for 5 min. Blocking buffer [0.4% Triton-100; 5% normal donkey serum in 1X PBS] was applied for 1 hr at room temperature. After blocking, primary antibodies were applied 1:1000 in blocking buffer (anti-MAP2 (Encor Biotechnology, Gainesville, FL, USA); anti-GFP, anti-RFP (Rockland Immunochemicals, Limerick, PA, USA)) and incubated overnight at 4 °C. The next day, three washes are performed with 0.1% Triton-100 in 1X PBS for 5 min each at room temperature. Secondaries were applied at 1:1000 (anti-chicken 647 (MAP2); anti-goat 488 (GFP); anti-rabbit 594 (RFP); Jackson ImmunoResearch, West Grove, PA, USA) and incubated for 2 hr at 4 °C. Slices/cells were then washed once with 0.1% Triton-100 in 1X PBS before DAPI (1:1000) was applied and incubated for 10 min, all at room temperature. Lastly, a final three washes for 5 min each in 0.1% Triton-100 in 1X PBS before mounting in Prolong gold anti-fade mounting media (ThermoFisher Scientific, Waltham, MA). All images were acquired on a Leica Stellaris 8 laser scanning confocal microscope (Leica Microsystems, Wetzlar, Germany), controlled by LAS X software.

### Cell culture

HEK293T cells were obtained from ATCC. Cells were maintained in DMEM medium (Gibco, ThermoFisher Scientific, Waltham, MA, USA) supplemented with 10% (v/v) FBS, penicillin (100 U mL^−1^), and streptomycin (100 µg mL^−1^) (ThermoFisher Scientific, Waltham, MA, USA) in a humidified incubator at 37 °C under 5% CO2. After transfection, cells were plated in DMEM containing 1% (v/v) dialyzed FBS, penicillin (100 U mL^−1^), and streptomycin (100 µg mL^−1^).

### Immunocytochemistry and co-immunoprecipitation of 5-HT_2A_ and mGluR2 in 5-HT_2A_ stable HEK cells

5-HT_2A_ stably expressing cells^5^ were transduced with mGluR2-mCherry lentivirus for two days. The day before experiments, cells were washed with PBS to remove 5-HT contained in 10% FBS and switched to 1% dialyzed FBS/DMEM. 5-HT_2A_ receptors were inducted with 1 µg/ml doxycycline for 18 hr prior to immunocytochemistry (ICC) or co-immunoprecipitation (co-IP) experiments. For ICC, cells were rinsed with 1X PBS and then incubated for 10 min with 4% PFA in PBS followed by 3x washes with 0.1%TX-100 in PBS. Cells were blocked for 30 min with 5% normal donkey serum/ 0.4% TX-100 in PBS and incubated at 4°C overnight with mouse anti-M1-FLAG (1:1000, #F3040; Sigma-Aldrich) and rabbit anti-RFP (1:1000, Rockland Immunochemicals, Limerick, PA, USA). The next day, cells were washed 3x with 0.1%TX-100 in PBS and with secondary antibodies incubation for 2 hr at room temperature (anti-mouse-Alexa 488 for M1-FLAG and anti-rabbit-Alexa594 for RFP, Jackson Immunoresearch). Cells were mounted and then imaged under a Leica STELLARIS 8 FALCON STED 100X objective.

For co-IP, cells were lysed in lysis buffer [1% N-dodecyl-β-*D*-maltoside (DDM), 20 mM Tris (pH7.4), and protease inhibitor (#11873580001; Sigma-Aldrich] and incubated for 1 hr at 4°C. The lysate was centrifuged for 10 min at 13,000 x g at 4°C and the supernatant was collected and incubated for 1 hr with FLAG-M2 beads (#M8823; Sigma-Aldrich). The flow-through fraction was collected after beads incubation. Beads were washed 3x with 0.5 mM NaCl, 20 mM Tris (pH 7.4). Proteins were eluted over 20 min with 2x Laemmli buffer at 55°C. Input, flow-through, and elution fractions were applied to 4-12% SDS-PAGE (#NP0336BOX; Life Technologies) and then transferred onto polyvinylidene fluoride (PVDF) membranes. Membranes were incubated with primary antibody, rabbit anti-FLAG (#F7425; SigmaAldrich) and rabbit anti-RFP (1:1000) antibodies overnight at 4°C followed by anti-rabbit-HRP secondary antibodies and finally reacted with enhanced chemiluminescence reagents. Images were captured with the ChemiDoc Image system.

### Bioluminescence resonance energy transfer (BRET2) assays

For mGluR2–5-HT_2A_ binding assays, human 5-HT_2A_, mGluR2, and mGluR3 were C-terminally fused to RLuc8 or GFP2 and subcloned into pcDNA3.1. For epitope-tagged constructs, a FLAG tag was placed at the N terminus. BRET2 experiments were performed based on established protocols^42,43^ with minor modifications. HEK293 cells were maintained in DMEM supplemented with 10% (v/v) FBS, penicillin (100 U mL−1), and streptomycin (100 µg mL−1) at 37 °C under 5% CO2. Cells were seeded at 1 × 10^4^ cells per well in poly-L-lysine-coated white 384-well plates and transfected 24 hr later using TransIT-2020 (Mirus; 3 µL per µg DNA). The total amount of DNA was held constant while varying the ratio of donor (RLuc8)- and acceptor (GFP2)-tagged receptors (GFP2/RLuc8 DNA ratio, 0–5). After an additional 24 hr, plates were backed with white film, culture medium was removed, and wells were washed once with 20 µL assay buffer (HBSS supplemented with 20 mM HEPES, pH 7.4, and 0.1% BSA). Assay buffer (20 µL) was added to each well, and GFP2 expression was verified by fluorescence measurement (485 nm excitation) using a PHERAstar FSX plate reader (BMG Labtech, Ortenberg, Germany). Coelenterazine-400a (NanoLight Technology/Prolume, Pinetop, AZ, USA) was then added (10 µL in assay buffer; 5 µM final concentration), and emissions at 395 nm (donor) and 510 nm (acceptor) were recorded using the PHERAstar FSX plate reader. BRET2 was calculated as the GFP2/RLuc8 emission ratio, and DNA ratio-dependent BRET2 data were analyzed in GraphPad Prism v10.

For concentration-response experiments, cells were transfected with 5-HT_2A_, mGluR2, GαoA–RLuc8, Gβ3, and Gγ8–GFP2 plasmids at a 1:1:1:1:1 ratio. When only one receptor was included, pcDNA3.1 was used to keep the total DNA amount constant. After an additional 24 hr, wells were washed with 20 µL per well of assay buffer, and 20 µL per well of assay buffer containing buffer alone, 10 µM LY379268, or 1 µM 25CN-NBOH was added. Plates were incubated for 15 min at 37 °C. Next, 10 µL of serial dilutions of 25CN-NBOH or LY379268 containing coelenterazine-400a (5 µM final concentration) was added, and plates were incubated at room temperature for 15 min before signals were detected as described above.

### GloSensor cAMP assay

Intracellular Gi-dependent inhibition of cAMP production was measured using the GloSensor-22F luciferase-based cAMP reporter (Promega). HEK293 cells were transiently transfected with GloSensor-22F together with human 5-HT_2A_, mGluR2, or both 5-HT_2A_ and mGluR2. Cells were seeded onto poly-L-lysine-coated white 384-well plates at 1 × 10^4 cells per well in Basal Medium Eagle (BME) without glutamate, supplemented with 1% dialyzed FBS.

After 24 hr, the medium was replaced with 20 µL per well of assay buffer (HBSS supplemented with 20 mM HEPES, pH 7.4, and 0.1% BSA) containing either buffer alone, 10 µM LY379268, or 1 µM 25CN-NBOH, and plates were incubated for 15 min at 37°C. Next, 10 µL of serial dilutions of 25CN-NBOH or LY379268 containing D-luciferin (GoldBio) at a final fixed concentration of 3 mM were added, and plates were incubated for 5 min at room temperature. Isoproterenol was then added (10 µL per well; 100 nM) to elevate endogenous cAMP via Gs. Luminescence was recorded after an additional 15 min incubation at room temperature using a SpectraMax L microplate luminometer (Molecular Devices).

### Nanobody-based protein purification and mass spectrometry

Two independent purification experiments were performed: one under basal conditions and one in which the selective 5-HT_2A_ agonist 25CN-NBOH was incorporated during the purification. For each experiment, whole brains were collected from 30 adult Htr2a-EGFP-CT mice (mixed male and female, postnatal days 30–60) via cervical dislocation and immediately snap-frozen in liquid nitrogen. Brains were stored at −80°C until use. On the day of purification, brains were placed in ice-cold homogenization buffer (TBS [20 mM Tris-HCl, pH 8.0, 150 mM NaCl], “cOmplete” Mini EDTA-free Protease Inhibitor Cocktail (Roche Diagnostics, Basel, Switzerland), 0.5 mM EDTA) and homogenized using a JOANLAB Overhead Stirrer Mixer (JOANLAB, Huzhou, China) at 1,500 rpm. For the agonist condition, 25CN-NBOH (10 µM final concentration) was included in the homogenization buffer and maintained at this concentration in all subsequent purification buffers throughout the experiment. The pooled homogenates were then divided equally into three aliquots (approximately 10 brains per aliquot) to serve as independent biological replicates (n=3 per condition). Crude brain homogenates were solubilized on a continuous rotor in 40 mM n-dodecyl-β-D-maltoside (DDM) for 1 hr at 4°C. Simultaneously, magnetic streptavidin beads (Promega, Madison, WI, USA) were washed with purification buffer [TBS; 0.5 mM DDM; 10 µM 25CN-NBOH for experimental condition], loaded with biotinylated nanobody construct for 30 min at 4°C, washed three times, pre-blocked with 0.5 mM biotin for 15 min, and washed three additional times with purification buffer. An anti-GFP nanobody was used for experimental samples and an anti-mCherry nanobody (LaM6) was used as a non-specific binding control, with identical purification conditions applied to both. Detergent-solubilized homogenates were clarified by centrifugation at 23,000 × g for 30 min, then each replicate was split in two equal volume fractions combined with nanobody-loaded beads (specific and non-specific nanobody loaded) and incubated on a rotor for 1 hr at 4°C. Beads were washed five times with 100 column volumes (CV) of purification buffer at 4°C using a magnetic rack. Site-specific elution was achieved by addition of 5 CV elution [purification buffer + 0.04 mg/mL SENP_EuB_ protease] followed by a 5-min incubation, with an additional 5 CV of purification buffer used to recover residual protein from the beads.

### Proteomics sample preparation

Immunoprecipitated samples were subjected to SDS-PAGE and stained with Coomassie blue. Lanes (1cm) for each sample were excised and cut into 1mm cubes and placed in 1ml of destain solution for 2 hr at room temperature (RT). After removal of destain solution, each sample was washed 2x with 100% Acetonitrile and incubated for 10 min at RT. Proteins were reduced with 10mM DTT for 10 min at 55 °C, then for 20 min at RT. After DTT was removed, samples were alkylated with 100mM IAA for 45 min in the dark at RT, and in-gel digested with 20ng/*μ*l of trypsin (Promega, Madison, WI, USA) overnight at 37°C. Peptides were extracted with 150 *μ*l of Acetonitrile, gently vortex and incubated at RT for 10 min. This was performed twice. Peptides were desalted with C18 spin columns (Pierce) and then dried via vacuum centrifugation. Peptide samples were stored at -80°C until further analysis.

### LC-MS/MS

The peptide samples were analyzed in technical duplicate by liquid chromatography-tandem mass spectrometry (LC-MS/MS) using an Ultimate3000 HPLC coupled to a Thermo OrbitrapExploris 480 mass spectrometer equipped with a Nanospray Flex ion source (ThermoFisher Scientific, Waltham, MA, USA). Peptides were eluted over an IonOpticks Aurora series 2 C18 column (75 *μ*m id × 15 cm, 1.6 *μ*m particle size; IonOpticks, Fitzroy, Victoria, Australia). Peptides were separated over a 120-min gradient at a 250 nl/min flow rate was used consisting of a 105-min linear gradient from 3-40% mobile phase B (MPB), and a constant 100% MBP for 15 min. Mobile phase A was 0.1% formic acid in water, and mobile phase B consisted of 0.1% formic acid in 80% acetonitrile. The Exploris480 was operated in Data Dependent Acquisition mode with a cycle time of 2s. Resolution for the precursor scan (m/z 375–1500) was set to 120,000, with AGC set to 300%. Following the full MS scan, a product ion scan was collected with a resolution set to 15,000, and normalized AGC set to 200%. The normalized collision energy was set to 30% for HCD. Peptide match was set to preferred, and precursors with unknown charge or a charge state of 1 and ≥ 7 were excluded.

### Data analysis

Raw data files were searched against the Uniprot reviewed mouse database (containing 17,230 entries, downloaded January 2025), the human HTR2A protein sequence, and a contaminants database, using the Sequest HT search engine node within Proteome Discoverer (v3.1, ThermoFisher Scientific, Waltham, MA, USA). Enzyme specificity was set to trypsin/P, up to two missed cleavage sites were allowed, methionine oxidation and N-terminus acetylation were set as variable modifications, and cysteine carbamidomethylation was set as a static modification. The Minora node was used to extract label-free quantification (LFQ) intensities. A 1% peptide-level false discovery rate (FDR) and a 5% protein-level FDR was used to filter all data. Match between runs was enabled. Mass spectrometry data have been deposited in the PRIDE repository under accession number PXD075336.

### Gene Ontology

Protein association networks and Gene Ontology (GO) biological process enrichment analyses were generated using the STRING database (v12). Proteins identified by mass spectrometry were uploaded to STRING using default parameters. Network edges represent known or predicted functional associations curated from existing databases, co-expression, text mining, and computational predictions, rather than direct physical interactions measured in this study. GO enrichment significance is reported as enrichment strength calculated by STRING.

### Western blot and immunoprecipitation in vivo

Whole brains from *Htr2a*^EGFP-CreERT2/EGFP-CreERT2^;*^Grm2^*^mCherry-Flpo/mCherry-Flpo^ mice were homogenized in lysis buffer [20 mM Tris, pH 7.4; 0.5 mM EDTA;150mM NaCl, 20mM DDM; “cOmplete” Mini EDTA-free Protease Inhibitor Cocktail] and incubated for 1 hr on a rotating mixer at 4°C. Tissue lysates were centrifuged at 15,000 × g for 30 min at 4°C. The supernatant (input) was collected and further incubated with GFP-Trap Magnetic Agarose beads (ChromoTek, Planegg-Martinsried, Germany) for at least 1 hr on a rotating mixer at 4°C. The flow-through fraction was collected after beads incubation. Beads were washed 3x with wash buffer (20mM Tris, pH7.4, 0.5mM EDTA, 150mM NaCl and 0.5mM DDM). Proteins on beads were eluted with 2X Laemmli buffer [4% SDS; 62.5 mM Tris, pH 7.4; 10% glycerol] at 55°C for 20 min. The input, flowthrough, and elution fractions were resolved by 4–12% SDS-PAGE (Life Technologies, ThermoFisher Scientific, Waltham, MA, USA) and transferred to PVDF membranes. Membranes were incubated overnight at 4°C with primary antibodies goat anti-GFP (1:1000) or rabbit anti-RFP (1:1000, Rockland Immunochemicals, Limerick, PA, USA), followed by horseradish peroxidase-conjugated anti-goat and anti-rabbit secondary antibodies (Jackson ImmunoResearch, West Grove, PA, USA). Membranes were finally reacted with enhanced chemiluminescence (ECL) reagents. Images were captured with the ChemiDoc Image system (Bio-Rad Laboratories, Hercules, CA, USA).

### Radioligand binding assays

#### Preparation of somatosensory cortex membrane fractions

Mice were euthanized by cervical dislocation and brains were rapidly removed. Somatosensory cortical tissue was dissected on ice and homogenized in ice-cold buffer [50 mM Tris-HCl, pH 7.4; 3 mM MgCl2; 1 mM EDTA; 1 mM DTT; “cOmplete” Mini EDTA-free Protease Inhibitor Cocktail]. Homogenates were centrifuged at 1,000 × g to pellet nuclei and debris (P1). The resulting supernatant (S1) was centrifuged at 25,000 × g for 20 min at 4°C. The pellet (P2) was resuspended in fresh buffer and centrifuged again; this wash step was repeated three times to remove endogenous ligands. Final pellets were resuspended in assay buffer [50 mM Tris-HCl, pH 7.4, containing Na⁺] and protein concentration was determined by Bradford assay (Bio-Rad Laboratories, Hercules, CA, USA) using BSA as a standard.

#### General conditions

Radioligand binding assays were performed using membrane preparations as described above. For competition binding assays, radioligands were used at concentrations approximating 0.5–1× Kd. Non-specific binding was defined using an excess of unlabeled ligand. All incubations were carried out for 60 min at 25°C unless otherwise indicated. Affinity (Kd) is defined as:

Kd=*k*off / *k*on

#### 5-HT2A receptor binding (DOI competition)

Competition binding assays were performed using [³H]ketanserin (∼0.5 nM) to label 5-HT_2A_ receptors. Membranes were incubated with radioligand and increasing concentrations of DOI in the absence or presence of the mGluR2/3 agonist LY379268 (10 µM). Reactions were terminated by rapid filtration through 0.3% polyethyleneimine (PEI)-soaked Whatman GF/B filters using a 48-channel Brandel harvester and washed with ice-cold 50 mM Tris-HCl (pH 7.4). Radioactivity retained on filters was measured by liquid scintillation counting in EcoScint A (National Diagnostics, 4 ml per filter). Radioligand competition binding data (retained dpm per filter) were analyzed using nonlinear regression implemented in Prism 10 (GraphPad Software): Data were fit using the built-in one-site (fit logIC50) and two-site competition ligand–receptor binding equations. Models were compared using extra sum-of-squares F test.

#### Kinetic [^125^I]DOI binding assays

Rat somatosensory cortex was prepared as described above and used to determine dissociation (k_off_) and association (k_obs_) rates. For k_off_, membranes were incubated with ∼100 pm [^125^I]DOI to equilibrium, at which point a saturating amount (final concentration: 10 µM) of cold DOI was added to each reaction at various times, after which the reactions were harvested onto PEI (0.3%)-soaked Whatman GF/B filters using a Brandel harvester and ice-cold wash buffer (50 mM Tris HCl, pH 7.4). Filter discs were dried overnight and counted in EcoScint A cocktail (4 ml/filter). Retained counts (dpm) were plotted as a function of time and fit to a one-phase exponential decay using GraphPad Prism 10 software. For k_obs_, [^125^I]DOI was added to membranes at various times, after which reactions were harvested and processed as described above, and data were fit to a one-phase exponential association curve using GraphPad Prism 10 software.

#### mGluR2/3 binding assays

Homologous competition binding assays were performed using [³H]LY341495 (∼1 nM) in membrane preparations. Membranes were incubated with radioligand and increasing concentrations of unlabeled LY341495. Data were analyzed using one-site and two-site competition models with F-test model selection. High- and low-affinity binding components were interpreted as corresponding to mGluR2 and mGluR3 sites, respectively.

#### Saturation binding (5-HT_2A_ receptor)

Radioligand saturation binding assays were performed using [³H]ketanserin across a range of concentrations in membrane fractions from wild-type and Grm2^−/−^ mice to determine Bmax (fmol/mg protein) and Kd.

#### Statistical analysis

All data are expressed as mean ± SEM, with statistical significance defined as p < 0.05. For behavioral studies, unpaired Welch’s t-tests were used to compare two groups. For single factor comparisons including Emax, BRETmax, and surface receptor expression, one-way ANOVA was performed followed by Sidak’s multiple comparison post hoc test. For experiments involving multiple groups or factors (RT-qPCR and cell surface ELISA), two-way ANOVA was performed followed by Dunnett’s or Sidak’s multiple comparisons post hoc test,respectively. Colocalization analysis for in vivo and in vitro imaging was performed using the JaCoP plugin in ImageJ and reported as Manders’ overlap coefficients (TM1 and TM2). BRET2 and GloSensor concentration-response data were analyzed by nonlinear regression using a three-parameter dose-response model. EC50, IC50, BRETmax, and BRET50 values were obtained from best fit parameters. Radioligand binding data were analyzed by nonlinear regression; one-site versus two-site models were compared using F-tests, and kinetic parameters were obtained by fitting exponential association and dissociation functions. All statistical analyses and curve fitting were performed in GraphPad Prism v10.

## Results

### Absence of direct 5-HT_2A_-mGluR2 interactions *in vitro* and *in situ*

Prior *in vitro* biochemical studies have reported contradictory results regarding direct interactions between 5-HT_2A_ and mGluR2, with some studies detecting no specific interactions^17^ and others reporting receptor multimers in transfected cells *in vitro*^21^. To investigate these competing hypotheses, we first performed immunocytochemistry and co-immunoprecipitation (Co-IP) studies with appropriately tagged receptors expressed in HEK293T cells to determine the potential for stable interactions *in vitro.* Our results show that both receptors are faithfully co-expressed on the cell surface (Fig. 1A), although mGluR2 was not detected in our co-IP studies with FLAG-tagged 5-HT_2A_ receptors (Fig. 1B) suggestive of an unstable interaction. To evaluate the potential for transient receptor-receptor interactions in living HEK293T cells, we utilized a saturation BRET2 assay and observed robust mGluR2-mGluR2 and mGluR3-mGluR2 interactions (Fig. 1C), consistent with the established homo- and heterodimerization of group II mGluR’s^44,45^. However, our 5-HT_2A_-mGluR2 BRET experiments revealed a signal equivalent to the negative control (*p*=0.8689), consistent with a lack of physical interaction (Fig. 1C and Supplemental Figure 1). These data suggest that 5-HT_2A_ and mGluR2 do not engage in direct protein-protein interactions, in agreement with Delille et al^17^.

**Figure 1.**
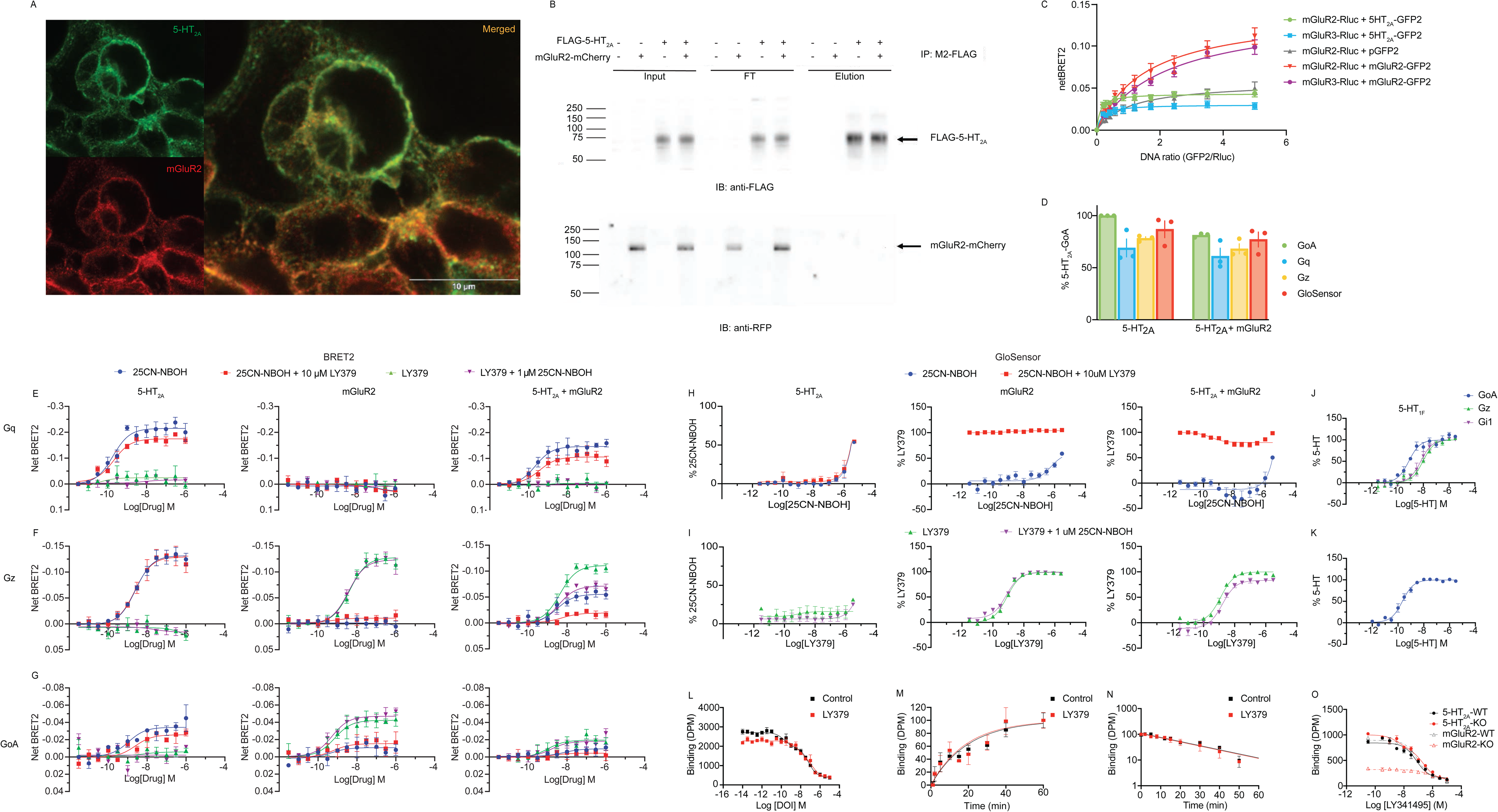
No receptor-receptor interactions in HEK293 cells. **(A)** Immunocytochemistry of mGluR2-mCherry and FLAG-5-HT_2A_ in HEK293T cells. Fixed with 4% paraformaldehyde 48 hours after transfection. Primary antibodies: anti-FLAG M1 antibody (1:1000) for FLAG-5-HT_2A_ and anti-RFP (1:1000) for mGluR2-mCherry. Secondary antibodies: anti-FLAG with anti-mouse Alexa647 (1:1000) and anti-RFP with anti-rabbit Alexa594 (1:1000). Images were acquisitioned under Leica STELLARIS 8 FALCON STED 100X/1.4 NA oil objective. **(B)** Cells prepared identically to (A) were harvested and cell lysates were used for pull-down with M2-FLAG antibody beads followed by input, flowthrough (FT) and elution fractions applied to SDS-PAGE for immunoblotting for FLAG-5-HT_2A_ against anti-rabbit FLAG antibody (1:1000) and mGluR2-mCherry fusion proteins against anti-RFP antibody (1:1000). **(C)** Receptor–receptor interactions measured by BRET2 saturation assays. RLuc8 tagged receptor was co-expressed with increasing amounts of GFP2-tagged receptor. Curves were fit to a one-site hyperbola model; BRET50 (acceptor:donor ratio at half-maximal BRET signal) are reported for each condition. mGluR2-RLuc8/mGluR2-GFP2: Bmax = 0.1406 ± 0.014, BRET50 = 1.605 ± 0.386; mGluR3-RLuc8/mGluR2-GFP2: Bmax = 0.141± 0.012, BRET50 = 2.188 ± 0.385; mGluR2-RLuc8/pGFP2: Bmax = 0.05734 ± 0.007, BRET50 = 1.052 ± 0.368; mGluR2-RLuc – 5-HT_2A_-GFP2: Bmax = 0.0435 ± 0.002, BRET50 = 0.1094 ± 0.034; mGluR3-RLuc – 5-HT_2A_-GFP2: Bmax = 0.03038 ± 0.002, BRET50 = 0.1648 ± 0.065. Data represent mean ± SEM of n = 3 independent experiments. **(D)** Surface co-expression of 5-HT_2A_ and mGluR2 receptors via ELISA assay. Receptor expression was detected using anti-FLAG HRP conjugated antibody (1:10,000) directed against the N-terminal FLAG-5HT_2A_ construct. Statistical comparisons were performed using two-way ANOVA followed by Sidak’s multiple comparisons post hoc test. Data are normalized to 5-HT_2A_-GoA condition and represent mean ± SEM of n = 3 independent experiments. **(E-G)** Net BRET2 measurements of Gαq (D), Gαz (E), and GαoA (F) coupling in HEK293T cells transfected with 5-HT_2A_ alone (left panels), mGluR2 alone (middle panels), or both 5-HT_2A_ and mGluR2 (right panels). Cells were treated with increasing concentrations of 25CN-NBOH (blue circles), 25CN-NBOH following 10 μM LY379268 (LY379) pretreatment (red squares), LY379 alone (green triangles), or LY379 following 1 μM 25CN-NBOH pretreatment (purple inverted triangles). Curves represent nonlinear regression fits to a three-parameter dose-response model. Data represent mean ± SEM of n = 3 independent experiments. **(H)** GloSensor cAMP assays measuring percent max 25CN-NBOH response for 5-HT_2A_ only (left panel) or percent maximal LY379 response for mGluR2 only (middle panel) and both 5-HT_2A_ and mGluR2 (right panel) from the 25CN-NBOH (blue circles) and 25CN-NBOH following 10 μM LY379 pretreatment (red squares) conditions. Data are normalized to the maximal response of the y-axis labeled reference agonist. Curves represent nonlinear regression fits to a three-parameter dose-response model. Data represent mean ± SEM of n = 3 independent experiments. **(I)** GloSensor cAMP assays measuring percent max 25CN-NBOH response for 5-HT_2A_ only (left panel) or percent maximal LY379 response for mGluR2 only (middle panel) and both 5-HT_2A_ and mGluR2 (right panel) from the LY379 alone (green triangles) and LY379 following 1 μM 25CN-NBOH pretreatment (purple inverted triangles) conditions. Data are normalized to the maximal response of the y-axis labeled reference agonist. Curves represent nonlinear regression fits to a three-parameter dose-response model. Data represent mean ± SEM of n = 3 independent experiments. **(J)** Net BRET2 measurements of GαoA (blue dots), Gαz (green triangles), and Gαi1 (blue dots) coupling in HEK293T cells transfected with 5-HT_1F_ alone. Cells were treated with increasing concentrations of 5-HT. Curves represent nonlinear regression fits to a three-parameter dose-response model. Data represent mean ± SEM of n = 3 independent experiments. **(K)** GloSensor cAMP assays measuring percent max 5-HT response for 5-HT_1F_. Data are normalized to the maximal response of the y-axis labeled reference agonist. Curves represent nonlinear regression fits to a three-parameter dose-response model. Data represent mean ± SEM of n = 3 independent experiments. **(L)** Competition radioligand binding of DOI at [^3^H]ketanserin-labeled 5-HT_2A_ sites in the presence or absence of mGluR2 agonist LY379 (1 μM). Binding is shown as mean ± SEM (DPM), with nonspecific binding fixed to the signal at the highest DOI concentration. Data were fit using one-site or two-site empirical models, with model selection based on *F*-tests and AIC. **(M)** Association time courses for [^125^I]DOI binding in cortical membranes under control conditions and in the presence of LY379. Data are mean ± SEM and were fit with a one-phase exponential association model. Outliers were identified using objective criteria and removed only when clearly aberrant. **(N)** Dissociation time course for [^125^I]DOI binding under control conditions and in the presence of LY379. Data are mean ± SEM (DPM) and were fit with a one-phase exponential model. **(O)** Homologous competition binding of [^3^H]LY341495 in membrane preparations from wild-type (WT) and knockout (KO) tissue for *Htr2a* (top) and *Grm2* (bottom). Binding is shown as mean ± SEM (DPM), with nonspecific binding constrained to the signal at the highest ligand concentration. Data were fit using one-site or two-site empirical models, with model selection based on *F*-tests and AIC. *Htr2*a datasets and *Grm2* WT were best described by two-site models, whereas *Grm2* KO was adequately fit by a one-site model.

Next, we explored claims that 5-HT_2A_ and mGluR2 interactions influence G-protein coupling selectivity *in vitro*. To evaluate these prior reports, we co-transfected various G-proteins (G_q_, G_z,_ G_oA_) in the presence of either 5-HT_2A_, mGluR2, or both 5-HT_2A_ and mGluR2 and determined signaling activity upon exposure to selective 5-HT_2A_ agonist 25CN-NBOH (10 *μ*M) and mGluR2/3 agonist LY379268 (10 *μ*M) with graded doses of the canonical agonist. To ensure co-expression of the mGluR2 and 5-HT_2A_ receptor did not alter surface receptor expression levels, surface ELISA assays were performed, revealing similar receptor expression levels across conditions (Fig. 1D; *F_(3,16)_*=0.3407, *p*=0.7962).

Confirming prior studies^46,47^, we observed robust activation of G_q_-heterotrimer dissociation via BRET studies at the 5-HT_2A_ receptor (Fig. 1E and Table 1). The maximum G_q_-BRET response (Supplemental Figure 2) was non-significantly diminished in cells co-transfected with mGluR2 and 5-HT_2A_ compared with 5-HT_2A_-expressing cells in all conditions except for 25CN-NBOH + LY379268-pretreated cells (Fig. 1E and Supplemental Figure 2). Agonist-stimulated G_z_ heterotrimer dissociation was significantly diminished in cells co-transfected with 5-HT_2A_ and mGluR2 compared with those transfected with 5-HT_2A_ alone following exposure to each agonist in their respective conditions (Fig. 1F and Supplemental Figure 2).

**Table 1.**
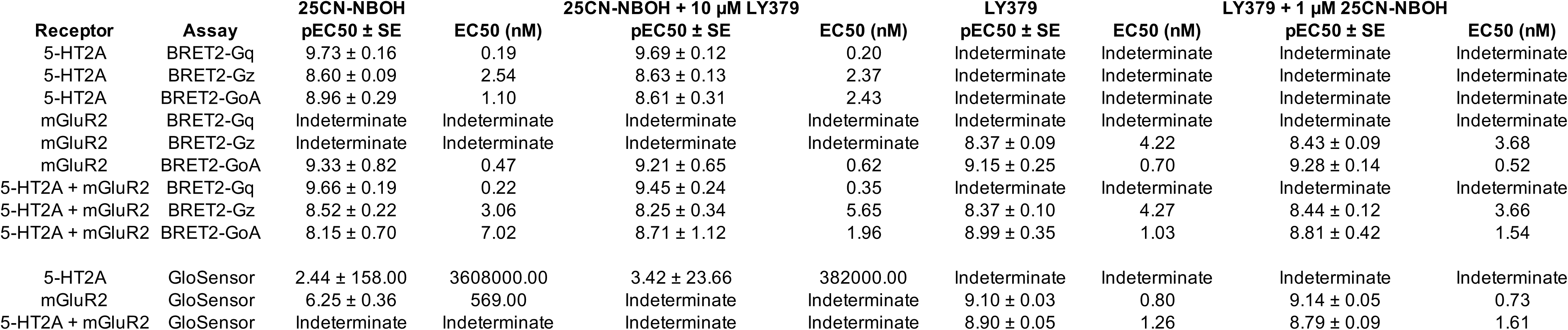
G-protein coupling BRET2 and GloSensor assay parameters. Responses were normalized to the maximal response of the reference agonist under the indicated assay condition. pEC50 ± standard error (SE) was calculated from three independent experiments, each performed in quadruplicate.

In cells co-expressing 5-HT_2A_ and mGluR2, no enhancement of G_oA_ BRET dissociation was observed as might be predicted based on prior studies^20,21,48^ (Fig. 1G and Supplemental Figure 2). Interestingly, we observed a small G_oA_ response in cells expressing 5-HT_2A_ alone following 25CN-NBOH treatment (Fig. 1G and Supplemental Figure 2) although a similar response was seen in the mGluR2 only condition (Fig. 1G and Supplemental Figure 2), suggesting coupling to endogenous receptor(s) rather than direct G_oA_ coupling to 5-HT_2A_. Canonical mGluR2-GoA signaling driven by addition of LY379268 (Fig. 1G and Table 1) was diminished in cells co-expressing 5-HT_2A_-mGluR2 (Fig. 1G and Table 1).

As an orthogonal measure, we tested the effects of treatment conditions employing direct measurements of cAMP inhibition between the three conditions. We observed a significantly increased cAMP inhibition in conditions where mGluR2 was co-transfected (Fig. 1H and 1I) (Supplemental Figure 3), indicating that inhibitory signaling response is due to the presence of mGluR2 and not 5-HT_2A_. This is consistent with the known G_i/o_ coupling of mGluR2^15,16^ and confirmed with control experiments where we found the 5-HT_1F_ (a canonical G_o/z/i_-coupled 5-HT receptors; Fig. 1J) robustly activated BRET dissociation and cAMP inhibition responses (Fig. 1K) which were not seen when cells were transfected with 5-HT_2A_ alone (Fig. 1H and 1I).

Prior studies revealed a putative allosteric effect of mGluR2 agonists on 5-HT_2A_ radioligand binding on membranes isolated from mouse cortex^21^. We attempted to replicate those previous findings which reported that activation of mGluR_2/3_ increases the high-affinity binding of hallucinogenic agonists at 5-HT_2A_ receptors in native cortical membranes. As originally described, the mGluR2 agonist LY379268 enhanced affinity of the agonist DOI for the 5-HT_2A_ receptor from 1.3 nM to 0.2 nM^21^. In our current studies, competition binding isotherms of DOI at [^3^H]ketanserin-labeled sites from WT mouse cortex were similar across conditions in the absence and presence of the mGluR2/3 agonist LY379268 (10 *μ*M) (Fig. 1L). Under control conditions, DOI competition was best described by a two-site model, consistent with the presence of high- and low-affinity binding components (Table 2. pIC_50,high_=9.69; pIC_50,low_=7.20; high-affinity fraction=0.33). Similarly, LY379268-treated data were described by a two-site model (Table 2. pIC_50,high_=8.29; pIC_50,low_=6.76). These data suggest that LY379268 does not significantly alter the two-affinity binding profile of DOI at 5-HT_2A_ receptors.

**Table 2.** Heterologous competition binding parameters for DOI at 5-HT2A receptors in the absence and presence of LY379268. Data from three independent experiments were analyzed by nonlinear regression using one-site and two-site ligand–receptor competition binding models. Values are presented as pIC50 ± standard error (SE), with corresponding IC50 values shown in nM. Extra sum-of-squares F-tests indicated that the two-site model was preferred for both groups: No LY379: F(2,71) = 33.18, p < 0.0001; LY379: F(2,63) = 5.119, p = 0.0087.

| Two Site competition | No LY379 |  | + LY379 |  |
| --- | --- | --- | --- | --- |
|  | pIC50 ± SE | IC50 (nM) | pIC50 ± SE | IC50 (nM) |
| pIC50-high | 9.69 ± 0.22 | 0.21 | 8.29 ± 0.43 | 5.17 |
| pIC50-low | 7.20 ± 0.12 | 62.62 | 6.76 ± 0.37 | 175.5 |

Because the reported effect of mGluR2 agonists is to enhance the affinity (K_d_) of DOI for 5-HT_2A_ receptors^21,22^, a direct prediction of this observation is that mGluR2 agonists should enhance agonist affinity by accelerating k_on_ or slowing k_off_. To test this possibility, we performed association and dissociation rate experiments employing [^125^I]DOI in the presence or absence of LY379268 in cortical membranes isolated from WT mice. We observed no effect of LY379268 on either association (Fig. 1M) or dissociation (Fig. 1N) kinetics of [^125^I]DOI on membrane preparations from mouse cortex. Dissociation rates (*k*_off_) were not significantly different in the absence and presence of LY379268 (Table 3; *k*_off_=0.037 ± 0.003 min^-1^ for both groups), and observed association rates (*k*_obs_) were likewise unchanged (Table 4; *k*_obs_=0.052 ± 0.005 min^-1^ for both groups), yielding similar calculated association rate constants (*k*_on_) under both conditions. Neither equilibrium nor kinetic analyses revealed a statistically significant effect of LY379268 on DOI binding to cortical 5-HT_2A_ receptors under the conditions of our assays. The absence of LY379268-dependent changes in both *k*_off_ and *k*_obs_ provides strong evidence against direct modulation of 5-HT_2A_ ligand binding by mGluR2.

**Table 3.** Dissociation kinetic parameters for [125I]DOI binding at 5-HT2A receptors. Dissociation kinetics of [125I]DOI binding were measured at 25°C using rat somatosensory cortex membrane preparations preincubated with 100 pM [125I]DOI in the absence or presence of LY379268 (LY379; 10 μM). Dissociation was initiated by addition of excess unlabeled DOI (10 μM). SB206553 (100 nM) was included throughout the assay to mask non-5-HT2A binding sites. Dissociation rate constants (k_off) were determined by fitting time-course data to a one-phase exponential decay model. Values are presented as k_off ± SE from three independent experiments. Fits with a shared k_off between groups and with unconstrained k_off values for each group were compared by extra sum-of-squares F test. A shared value for k_off was preferred: F(1,83) = 0.1005, p = 0.1005.

| Dissociation kinetic parameter | No LY379 | + LY379 |
| --- | --- | --- |
| | $k_{\text{off}} \pm \text{SE (min}^{-1}\text{)}$ | $k_{\text{off}} \pm \text{SE (min}^{-1}\text{)}$ |
| | $0.037 \pm 0.003$ | $0.037 \pm 0.003$ |

**Table 4.** Association kinetic parameters for [125I]DOI binding at 5-HT2A receptors. Association kinetics of [125I]DOI binding were measured at 25°C using rat somatosensory cortex membrane preparations incubated with 100 pM [125I]DOI in the absence or presence of LY379268 (LY379; 10 μM). SB206553 (100 nM) was included throughout to mask non-5-HT2A binding sites. Observed association rate constants (k_obs) were determined by fitting time-course data to a one-phase exponential association model. Values are presented as k_obs ± SE from three independent experiments. Fits with a shared k_obs between groups and with unconstrained k_obs values for each group were compared by extra sum-of-squares F test. A shared value for k_obs was preferred: F(1,65) = 0.0828, p = 0.7744.

| Association kinetic parameter | No LY379 | + LY379 |
| --- | --- | --- |
| | $k_{\text{obs}} \pm \text{SE (min}^{-1}\text{)}$ | $k_{\text{obs}} \pm \text{SE (min}^{-1}\text{)}$ |
| | $0.052 \pm 0.006$ | $0.052 \pm 0.006$ |

To determine whether the observed binding components depend on 5-HT_2A_ receptor expression, we performed homologous competition binding assays using [^3^H]LY341495 in cortical membrane preparations from wild-type, *Htr2a* knockout^40^, and *Grm2* knockout mice^39^. In wild-type and *Htr2a* knockout membranes, LY341495 binding was best described by a two-site model, indicating the presence of multiple binding components. In contrast, *Grm2* knockout membranes were adequately described by a one-site model, consistent with loss of a distinct binding component. Global comparison of two-site models across wild-type and *Htr2a* knockout datasets revealed no significant differences in binding parameters (Fig. 1O and Table 5). These results indicate that the biphasic binding profile of LY341495 depends on mGluR2 expression but does not require 5-HT_2A_ receptor expression. Together, these findings indicate that mGluR2 agonists do not modulate 5-HT_2A_ ligand binding under either equilibrium or transient (*e.g.*, kinetic) conditions.

**Table 5.** Homologous competition binding parameters for [3H]LY341495 binding at mGluR2/3 receptors. Homologous competition binding assays were performed using ∼1 nM [3H]LY341495 in rat somatosensory cortex membrane preparations from 5-HT2A knockout, mGluR2 knockout, and corresponding wild-type animals. Membranes were incubated with radioligand and increasing concentrations of unlabeled LY341495. Data from three independent experiments were fitted by nonlinear regression using one- and two-site ligand–receptor competition binding models. Values are presented as pIC50 ± SE, with corresponding IC50 values shown in nM. Extra sum-of-squares F-tests indicated that the two-site model was preferred for all groups except mGluR2-KO: 5-HT2A-WT: F(2,17) = 7.489, p = 0.0047; 5-HT2A-KO: F(2,19) = 8.416, p = 0.0024; mGluR2-WT: F(2,19) = 10.65, p = 0.0008; mGluR2-KO: F(2,18) = 2.103, p = 0.1511.

|  |  |  |  |  |  |  |  |  |
| --- | --- | --- | --- | --- | --- | --- | --- | --- |
| One Site competition |  |  |  |  |  |  | mGluR2-KO<br>pIC50 ± SE<br>6.30 ± 0.22 | IC50 (nM)<br>498.3 |
| Two Site competition |  | 5-HT2A-WT<br>pIC50 ± SE | IC50 (nM) | 5-HT2A-KO<br>pIC50 ± SE | IC50 (nM) | mGluR2-WT<br>pIC50 ± SE | IC50 (nM) |  |
| pIC50-high |  | 8.39 ± 0.38 | 4.04 | 9.12 ± 0.41 | 0.76 | 8.70 ± 0.35 | 2.02 |  |
| pIC50-low |  | 6.66 ± 0.20 | 220.6 | 6.81 ± 0.12 | 155.4 | 6.72 ± 0.11 | 192.6 |  |

### Proteomic analysis of brain 5-HT_2A_ receptor interacting proteins

We next attempted co-immunoprecipitation (Co-IP) studies from mouse cortex but were hampered by the lack of specific and suitably sensitive antibodies for either the 5-HT_2A_ or mGluR2 receptors. Fortunately, we recently created GFP-tagged Htr2a-expressing mice (Htr2a-EGFP-CT mice; Chiu et al., *in press)* where we CRISPR-tagged endogenous 5-HT_2A_ receptors at the genomic locus. As previously shown by us^49–51^ and others^52–54^ an intact PDZ-domain interaction motif (e.g. SCV in the C-terminus) is essential for the function and appropriate cellular and subcellular localization of 5-HT_2A_ receptors. Importantly, the expression, distribution and function of Htr2a-EGFP-CT is essentially equivalent to WT *in vivo* (Chiu et al., *in press).* With these mice in hand, we next developed receptor purification workflow based on a recently reported nanobody-based system^55^. We refer to this approach as “RAPID” (**R**eceptor **A**ffinity **P**urification **I**n One **D**ay^14^) which enables native 5-HT_2A_-EGFP receptor isolation from mouse brain tissue in ∼3 hr for proteomics experiments (Fig. 2A).

**Fig 2.**
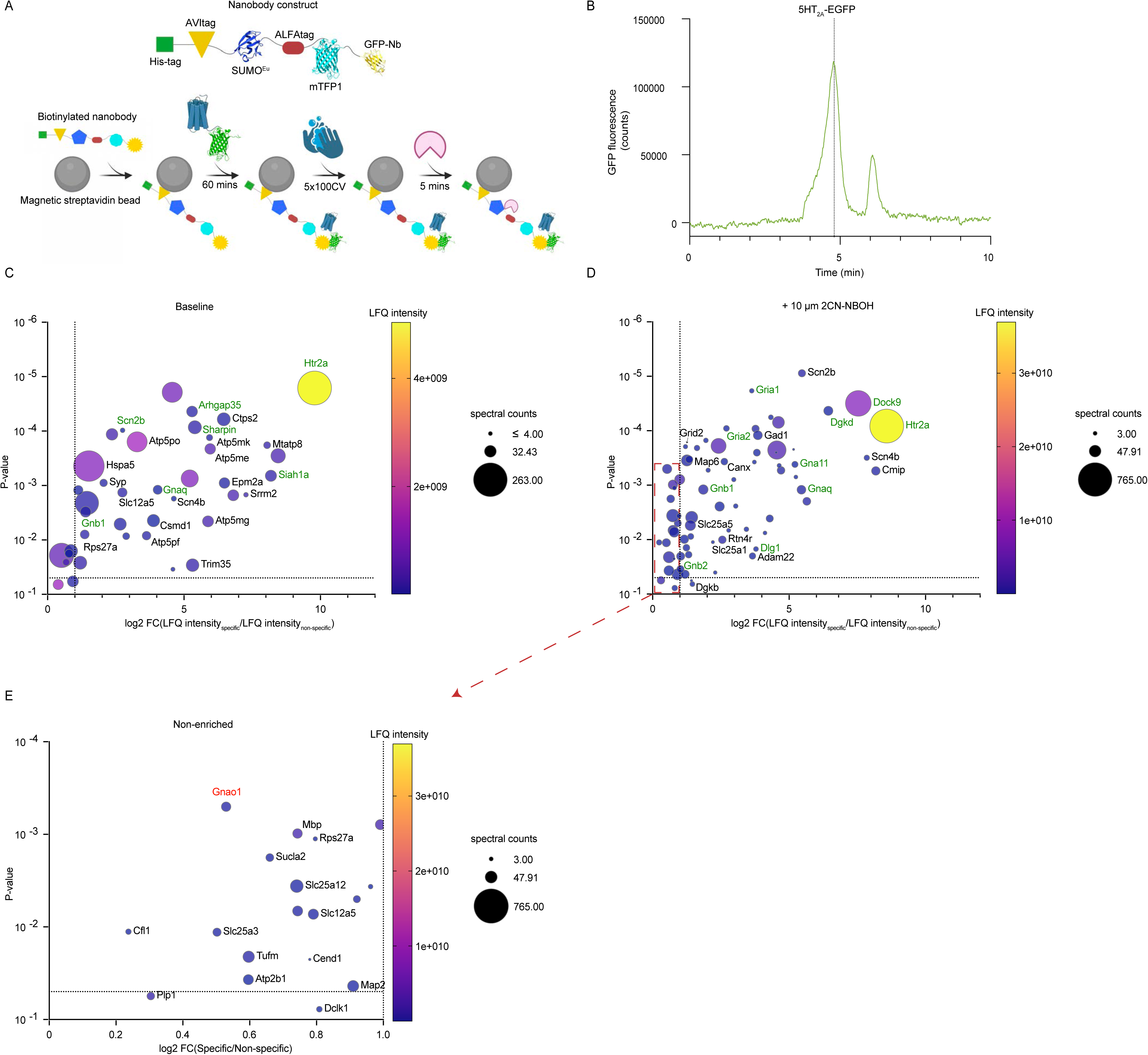
RAPID purification of 5-HT_2A_–EGFP-CT reveals interactions with synaptic proteins. **(A)** Schematic of the Receptor Affinity Purification In One Day (RAPID) workflow used to isolate native 5-HT_2A_–EGFP-CT from brain tissue. **(B)** Fluorescence-detected HPLC trace confirming elution of stable purified 5-HT_2A_–EGFP-CT from brain tissue of Htr2a-EGFP-CT mice. **(C)** Volcano plot of mass spectrometry results comparing native 5-HT_2A_–EGFP-CT purified via GFP nanobody to a non-specific LaM6 (mCherry) nanobody control under basal (vehicle) conditions. The x-axis represents log2 fold-enrichment; the y-axis represents -log10(p-value). Yellow points indicate the most abundant protein enriched in the sample. Green gene labels indicate proteins of interest (Htr2a, Arhgap35, Sharpin, Siah1a, Scn2b, Gnaq, Gnb1). Selection criteria were proteins enriched greater than or equal to 1.2-fold-change and with a p-value < 0.05. **(D)** Volcano plot as in (C) from samples in which 10 μM 25CN-NBOH was present throughout purification. Yellow points indicate the most abundant protein enriched in the sample. Green gene labels indicate proteins of interest (Htr2a, Dock9, Dgkd, Scn2b, Scn4b, Gnaq, Gna11, Gria1, Gria2, Gnb1, Dlg1). Selection criteria were proteins enriched greater than or equal to 1.2-fold-change and with a p-value < 0.05. Dotted red box correlates to the proteins shown in (E). **(E)** Enlarged view of the non-enriched proteins from the red box in (D).

Whole brains were solubilized with mild maltoside detergents to preserve native interactions present in the tissue. The detergent solubilized brain lysates were incubated with a GFP nanobody construct to selectively isolate 5-HT_2A_-receptor interacting proteins from our Htr2a-EGFP-CT mice, using an unrelated nanobody construct (mCherry; LaM6^56^) as a non-specific control. Purified complexes were monodisperse as assessed by GFP fluorescence-detection high-pressure liquid chromatography (FLD-HPLC), confirming isolation of biochemically stable receptor complexes (Fig. 2B). Mass spectrometry analysis revealed that 5-HT_2A_ was the most enriched protein with a log_2_ fold change of 9.93, corresponding to a 978-fold increase compared to the non-specific control, under basal conditions (Fig. 2C). Similar studies were then performed in the presence of the highly selective psychedelic 25CN-NBOH 5-HT_2A_ agonist (Figs. 2D, 2E). Although we did not detect mGluR2 protein, known 5-HT_2A_ interacting (e.g. *Dlg1*/SAP97) and canonical signaling partners (e.g. *Gnaq*/Gα_q_, Gna11/Gα_11_, Gnb1/Gβ_1_ and Dgkd/Diacylglycerol Kinase Delta) were readily identified. In the agonist-treated condition, the G-protein subunits *Gnaq* and *Gna11* (Gα_q_ and Gα_11_) were selectively enriched relative to control (Fig. 2D), while other G-protein subunits were either not enriched (Gnao1/Gα_oA_, *Gnaz*/Gα_z,_ and Gnb2/Gβ_2_) or were undetected (Gnai1/Gα_i1_; Fig. 2E).

Mass spectrometry detected 531 proteins across both conditions. Applying a false discovery rate (FDR) threshold of ≤0.05 and a log2 fold-change cutoff of ±0.26 (∼1.2-fold), 23 proteins were enriched in samples purified in the presence of 25CN-NBOH relative to control. Examining both datasets allowed us to identify enriched proteins unique to the agonist-exposed dataset as well as those shared by both (Fig. 3A). STRING database GO analysis^57^ of the proteins enriched in the 25CN-NBOH group revealed selective enrichment in glutamatergic synapses (Figs. 3B, 3C), consistent with prior reports that 5-HT_2A_ receptors are preferentially localized to Layer V glutamatergic neurons^25–28,38,58,59^. Specific enrichment was also identified for proteins localized to post-synaptic densities (PSDs) and synapses (Fig. 3C). These results are consistent with many prior studies that 5-HT_2A_ receptors interact with PDZ-domain proteins like *DLG1* (SAP97) which are localized to excitatory PSDs^49–53,60–65^. Significantly, analysis of proteins common to both datasets identified a cellular context describing G protein signaling, potentially reflecting receptor-G protein interactions under basal conditions (Figs. 3D, 3E).

**Fig 3.**
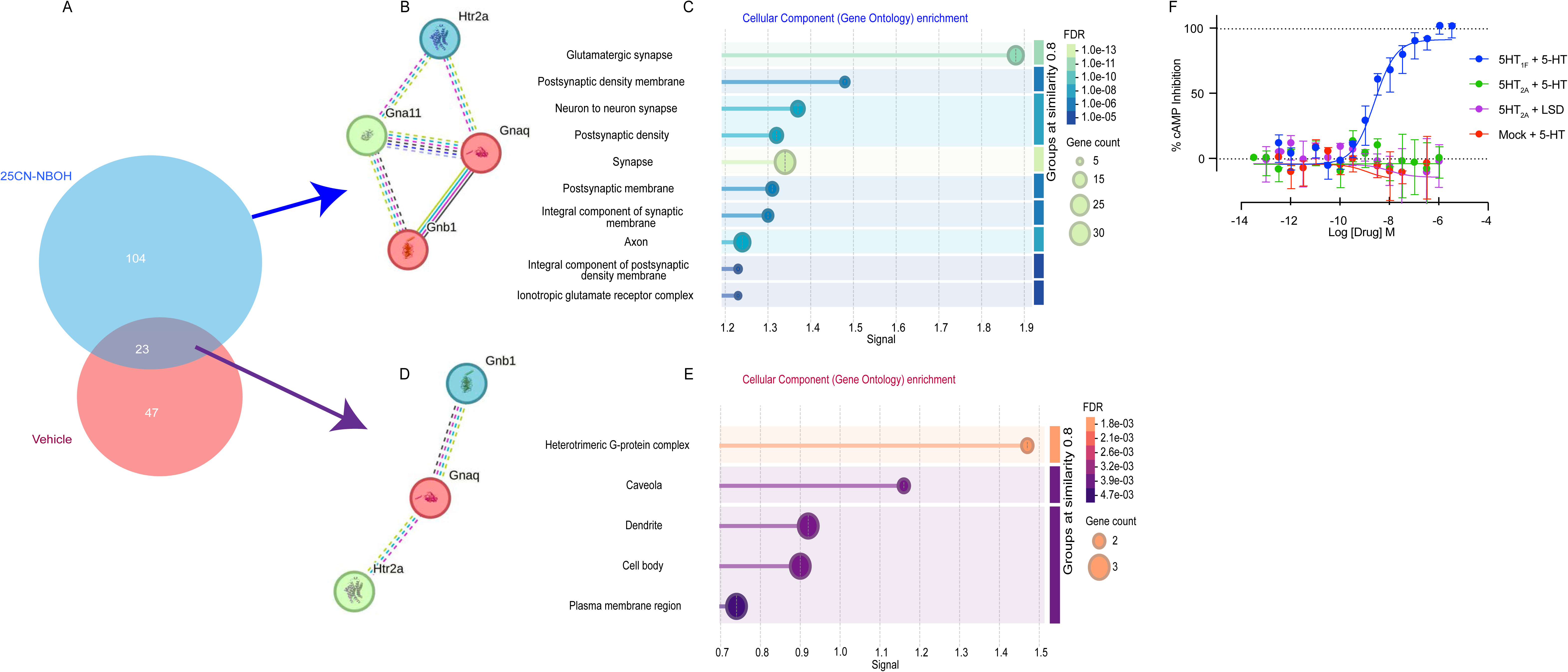
Protein interaction network analysis of 5-HT_2A_ Interacting proteins. **(A)** Venn diagram (DeepVenn) of proteins enriched in 5-HT_2A_-EGFP-CT pulldowns under basal (vehicle; red; n = 47 unique proteins) and agonist-treated (25CN-NBOH; blue; n=104 unique proteins) conditions. Purple indicates proteins identified under both conditions (n = 23). **(B)** STRING protein-protein interaction network of all enriched 5-HT_2A_ interacting proteins. K-means clustering (k = 3) was applied; node colors represent cluster assignment. Minimum interaction score = 0.400 (medium confidence). **(C)** Cellular Component Gene Ontology enrichment analysis (STRING.org) corresponding to the clusters shown in (B). FDR threshold of < 0.05. **(D)** STRING interaction network restricted to the proteins identified under both basal and 25CN-NBOH conditions (purple overlap from A), with k-means clustering (k = 3). Minimum interaction score = 0.400 (medium confidence). **(E)** Cellular Component Gene Ontology enrichment analysis for the proteins in (D). **(F)** G_i_ -mediate isoproterenol-stimulated cAMP accumulation in HEK293 cells expressing 5-HT_1F_ (blue), 5-HT_2A_ (green), or untransfected mock cells (red) in the presence of 5-HT (10μM) and 5-HT_2A_ in the presence of LSD (10μM, purple). Data represent mean ± SEM; n = 3 independent experiments.

Together, these findings do not support a recent proposal that psychedelic drugs utilize Gα_i_ for signaling *in vitro* and *in vivo* for their psychedelic actions^66^. To further examine the proposal that 5-HT_2A_ receptors can productively couple to Gα_i_, we examined the ability of the psychedelic drug lysergic acid diethylamide (LSD) and the non-psychedelic agonist 5-HT to inhibit agonist-stimulated cAMP production—the canonical action of Gi-coupled GPCRs. As shown in figure 3F, activation of the 5-HT_1F_ serotonin receptor (which canonically couples to Gα_i_) inhibited isoproterenol-stimulated cAMP production while activation of 5-HT_2A_ receptors by either the full agonist 5-HT or the psychedelic LSD had no effect. Collectively, these results are consistent with prior studies demonstrating that 5-HT_2A_ receptors couple to Gα_q/_Gα_11_ *in vivo*^13,46,67,68^.

### Creation and characterization of double knock-in Htr2a-EGFP-CT x mGluR2-mCherry mice

As proteomics studies suffer from the inherent liability that weak or transient interactions will not be stabilized during detergent purification, we re-evaluated the possibility that 5-HT_2A_ and mGluR2 receptors interact *in situ* in neurons. Because suitably selective and sensitive antibodies for mGluR2 (or 5-HT_2A_) receptors do not exist, we created a mouse in which the endogenous mGluR2 locus was CRISPR-tagged with the fluorescent protein mCherry, as has been done for other GPCR mapping studies^38,69^. We verified the functionality of mGluR2-mCherry fusion protein in transfected HEK-293 cells and found equivalent activity compared with the WT receptor^14^. We then visualized mGluR2 tagged receptors by confocal microscopy in our Grm2-mCherry-CT-IRES-Flpo mice (Fig. 4A) which display a distribution similar to that previously reported for mGluR2^23,37^.

**Fig 4.**
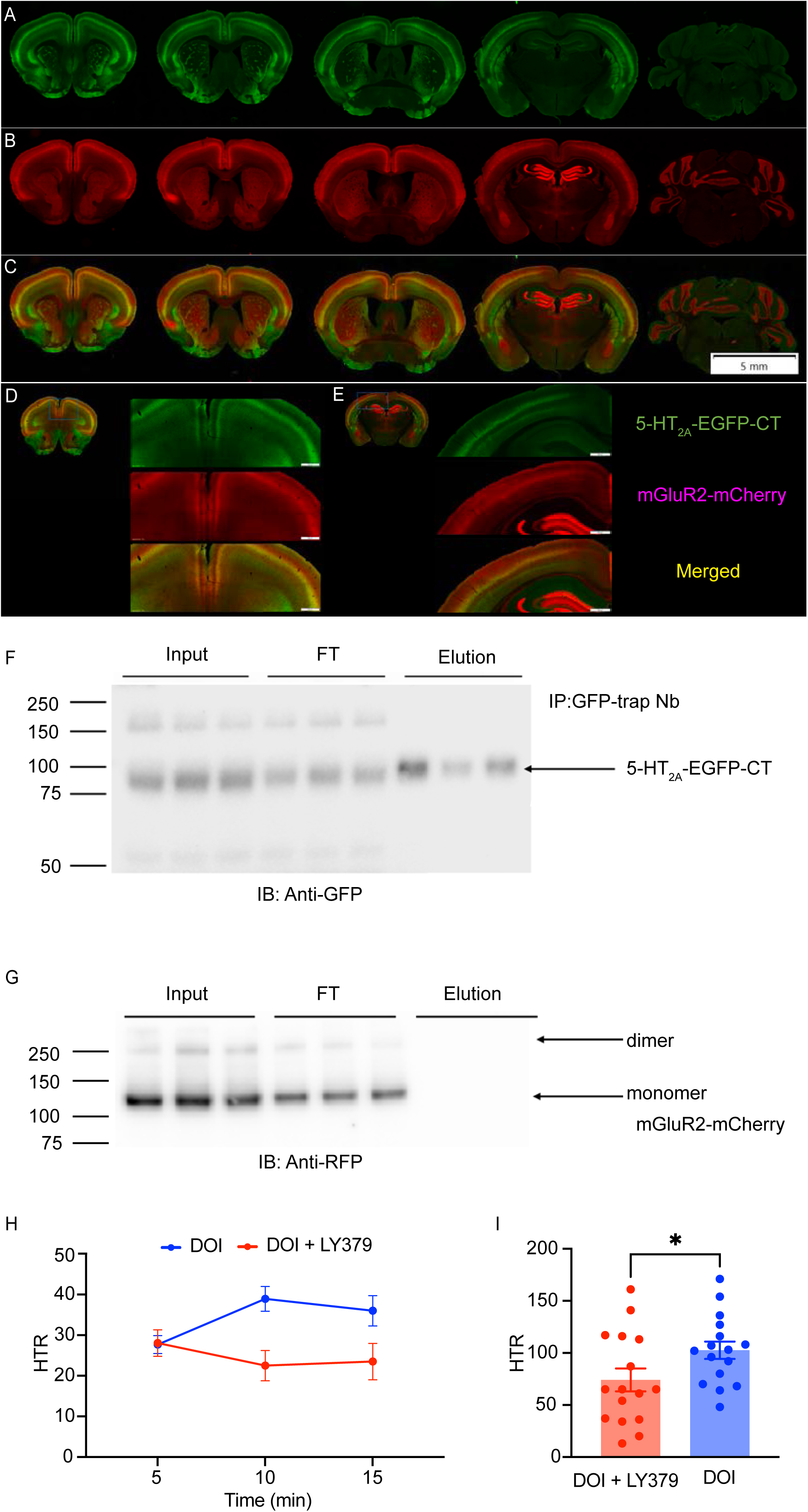
5-HT_2A_ and mGluR2 expression patterns throughout the mouse brain. **(A)** Representative coronal brain slices from adult Htr2a-EGFP-CT × mGluR2-mCherry^+/+^ mouse showing 5-HT_2A_-EGFP-CT (488 nm) **(B)** mGluR2-mCherry (594 nm) and **(C)** merged images. Primary antibodies: anti-GFP antibody (1:1000) for 5-HT_2A_-EGFP and anti-RFP (1:1000) for mGluR2-mCherry. Secondary antibodies: anti-goat Alexa488 with anti-rabbit Alexa594 (1:1000) and anti-RFP with anti-rabbit Alexa594 (1:1000). All images were acquired with a 10X objective using a VS200 slide scanner. Scale bar = 2 mm. **(D)** Representative anterior coronal section from an adult Htr2a-EGFP-CT × mGluR2-mCherry^+/+^ mouse. Top left corner: merged overview of the full brain section. Zoomed panels show cortical regions: 5-HT_2A_-EGFP-CT (top), mGluR2-mCherry (middle), and merged (bottom). All images were acquired with a 10X objective using a VS200 slide scanner. Scale bar = 500 μm. **(E)** Representative hippocampal coronal section from the same Htr2a-EGFP-CT × mGluR2-mCherry^+/+^ mouse. Top left: merged overview. Zoomed panels show cortical and hippocampal regions: 5-HT_2A_-EGFP-CT (top), mGluR2-mCherry (middle), and merged (bottom). All images were acquired with a 10X objective using a VS200 slide scanner. Scale bar = 500 μm. **(F)** Co-immunoprecipitation from cortex tissue lysates of three adult Htr2a-EGFP-CT × mGluR2-mCherry^+/+^ mice. GFP-tagged receptor was immunoprecipitated using GFP nanobody; blots were probed with anti-GFP antibody to detect 5-HT_2A_. Input, flow through (FT) from lysate, and elution were all used for western blot results. **(G)** Co-immunoprecipitation as in (C); blots probed with anti-RFP antibody to detect mGluR2. **(H)** Time course of head-twitch responses (HTR) in adult Htr2a-EGFP-CT × mGluR2-mCherry^+/+^ mice following treatment with DOI (1 mg/kg) (blue) or DOI proceeded by 30-min pretreatment with mGluR2 agonist LY379268 (LY379; 15 mg/kg; red). Data represent mean ± SEM; n = 16 per group. **(I)** Total HTR counts over a 15-minute window for groups shown in (E). (DOI = 102.6 ± 8.331; DOI + pretreatment = 74.06 ± 11.03 head twitches per 15 min; p = 0.0482, Welch’s t-test; n=16 per group).

To create a double knock-in strain containing fluorescent markers on the two proteins of interest, we crossed the Grm2-mCherry-CT-IRES-Flpo with our previously described Htr2a-EGFP-CT-IRES-CreERT2 mouse line^38^. Initial mapping studies in these heterozygous double knock-in mice revealed that both 5-HT_2A_ and mGluR2 receptors are localized to discrete cortical layers with potential overlapping distributions (Figs. 4A-C). Clear laminar patterning of the two receptor types is observed, with mGluR2 enriched in layer IV and V and 5-HT_2A_ enriched in layer Va (Figs. 4C-E). We found mGluR2 receptors were expressed at relatively low levels in cortical layers IV and V, with lower levels throughout the neuropil of several cortical layers. In contrast to a similar distribution in multiple cortical areas, significant differences in expression were noted in other brain areas. Thus, mGluR2 receptors were most highly expressed in the hippocampal subfields (CA1, CA3, and dentate gyrus) and cerebellum (Figs. 4A, 4C), consistent with previous findings^23,37^. 5-HT_2A_ receptors, by contrast, are expressed at low levels in hippocampus and were not detected in the cerebellum (Figs. 4B, 4C). In the striatum, 5-HT_2A_ receptors show a patch-like distribution while mGluR2 display a low, matrix-like expression pattern^70^. Using cortical brain tissue from the Htr2a-EGFP-CT x mGluR2-mCherry double knock-in mice, we performed co-immunoprecipitation studies using a commercial GFP nanobody to isolate 5-HT_2A_-EGFP fusion protein (Fig. 4F). Both monomer and dimer mGluR2-mCherry were detected in the lysate input and flow through (FT) but not in the elution fractions (Fig. 4G), suggesting that the two receptors do not form stable complexes in mouse brain tissue.

We then used these mice for behavioral, biochemical and anatomical studies to thoroughly test the predictions arising from the competing hypotheses regarding mGluR2 modulation of psychedelic drug actions. We replicated prior findings, using our double knock-in Htr2a-EGFP-CT x mGluR2-mCherry mice, that pre-treatment with an mGluR2 agonist reduces DOI-induced HTR^21,30,31^. As shown, pretreatment with the mGluR2 agonist LY379268 (15 mg/kg) attenuates the DOI-induced HTR in these double knock-in mice compared to saline pretreatment (Figs. 4H, 4I; *t*(27.91)=2.066, *p*=0.0482).

#### 5-HT_2A_ and mGluR2 receptors do not co-localize *in vivo*

Immunofluorescent imaging studies using confocal microscopy revealed a pattern of 5-HT_2A_ expression identical to that previously reported for native 5-HT_2A_ receptors^25,26,36^. Specifically, 5-HT_2A_ receptors are localized to the layer V pyramidal neuronal somatodendritic compartment with apical dendrites expressing 5-HT_2A_ receptors projecting to layer I (Fig. 5A).

**Fig 5.**
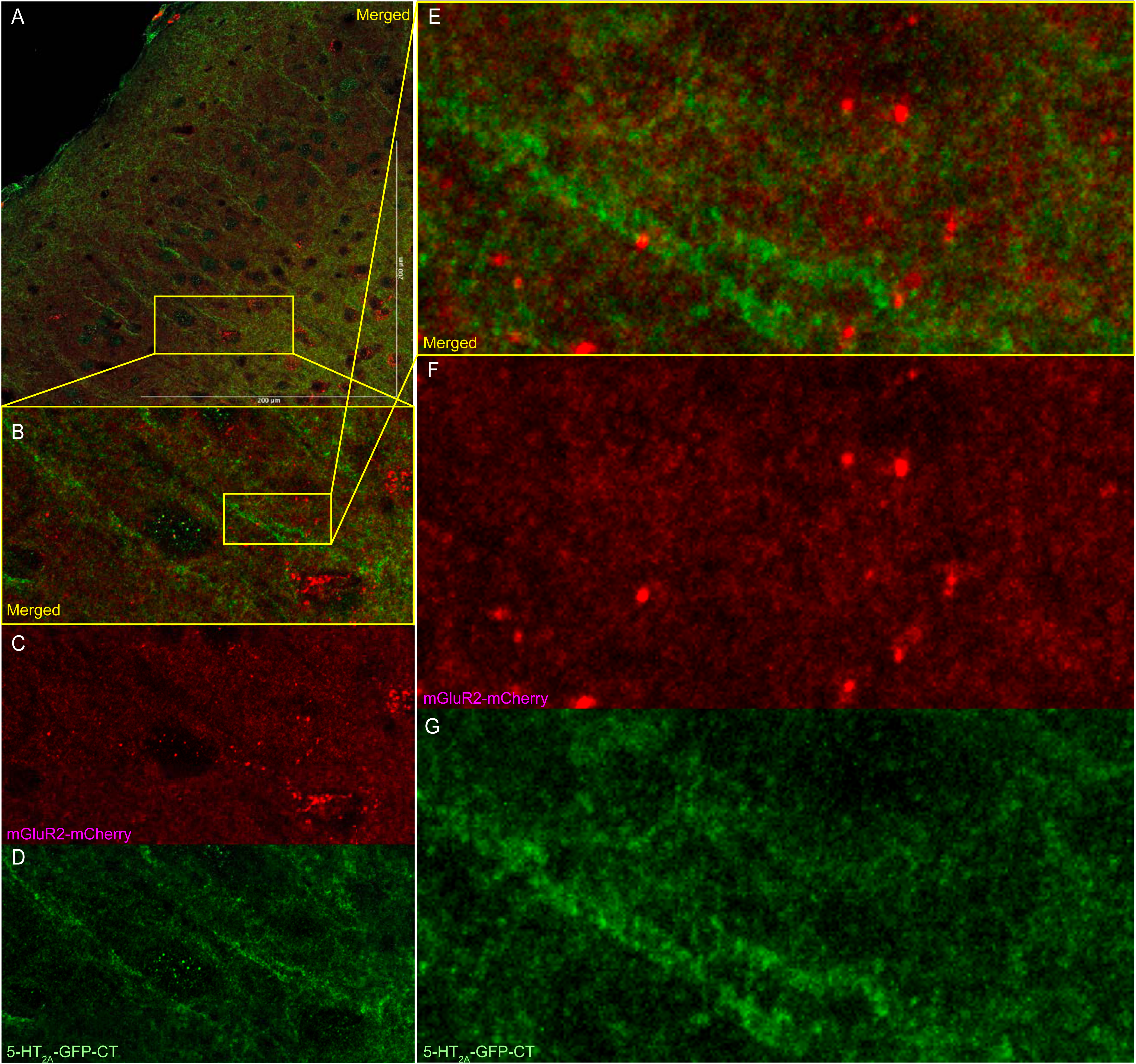
5-HT_2A_ and mGluR2 receptors do not co-localize in somatosensory cortex. **(A)** Representative confocal images of somatosensory cortex (S1) from an adult Htr2a-EGFP-CT × mGluR2-mCherry^+/+^ mouse, showing 5-HT_2A_-expressing layer 5 pyramidal neuron apical dendrites projecting through the superficial cortical layers. **(B, E)** Zoomed views from (A). **(C, F)** mGluR2-mCherry (RFP) channel only, corresponding to views in (B) and (E), respectively. **(D, G)** 5-HT_2A_-EGFP-CT (GFP) channel only, corresponding to views in (B) and (E), respectively. All images were acquired using a Leica STELLARIS 8 FALCON STED 100X/1.4 NA oil objective and 4X frame averaging with 1X zoom.

We closely examined the potential for mGluR2 and 5-HT_2A_ receptor co-localization in the primary somatosensory cortical region (S1). 5-HT_2A_ receptors were localized to apical dendrites (Figs. 5B, 5D, 5G) while mGluR2 receptors were found in scattered puncta throughout the cortex as well as diffusely in the neuropil (Figs. 5C, 5F). Minimal co-localization of Htr2a-EGFP-CT and mGluR2-mCherry was observed through analysis of Mander’s coefficients (Supplemental Figure 4, TM1=0.283; TM2=0.172). Here, TM1 reflects the fraction of GFP signal overlapping mCherry, whereas TM2 reflects the mCherry fraction overlapping GFP; a Mander’s coefficient of 1 indicates complete overlap. These results suggest little to no overlap occurs between these two channels *in vivo*.

#### 5-HT_2A_ and mGluR2 receptors do not co-localize in neurons *in vitro*

We next evaluated the potential co-localization of 5-HT_2A_ and mGluR2 receptors in primary cortical neurons *in vitro* as recent studies have suggested a translation-independent association of mRNAs that encode protomers of the 5-HT_2A_-mGluR2 receptor complex^71^. Specifically, Htr2a and Grm2 transcripts reportedly interact physically in vitro and in brain tissue, potentially facilitating coordinated receptor expression that forms the proposed 5-HT_2A_-mGluR2 heterodimer complex.

Prior studies have shown mGluR2 expression peaks at post-natal day 14 (P14) reaching mature levels at P21^72^ with mGluR2 binding sites gradually increasing in cortical layer II, reaching adult density at P21. By contrast, 5-HT_2A_ mRNA levels appear to peak around P5 whereas the number of binding sites peak at P13^73^. We first use rt-qPCR to quantify the expression of 5-HT_2A_ (*Htr2a*) and mGluR2 (*Grm2*) mRNAs in cortical neurons *in vitro* at various days in vitro (DIV) derived from double knock-in Htr2a-EGFP-CT x mGluR2-mCherry mice or wild type (WT) C57BL/6J mice (Fig. 6A). Peak mRNA expression for both genes occurred at DIV14 (WT, *Htr2a*, *F_(4,10)_*=20.93, p<0.0001; WT, *Grm2*, *F_(4,10)_*=24.89, p<0.0001; 5-HT_2A_xmGluR2, *Htr2a*, *F_(4,10)_*=26.73, *p*<0.0001; 5-HT_2A_xmGluR2, *Grm2*, *F_(4,10)_*=14.66, *p*=0.0003), corresponding roughly to the second postnatal week in mice. Lower mRNA levels observed at DIV21 may reflect increased protein accumulation as neurons reach developmental maturity rather than reduced transcriptional activity. To verify these findings at the protein level in vivo, immunohistochemistry of brain tissue at P0 (Fig. 6B) and P30 (Fig. 6C) reveal an apparent increase in receptor expression.

**Fig 6.**
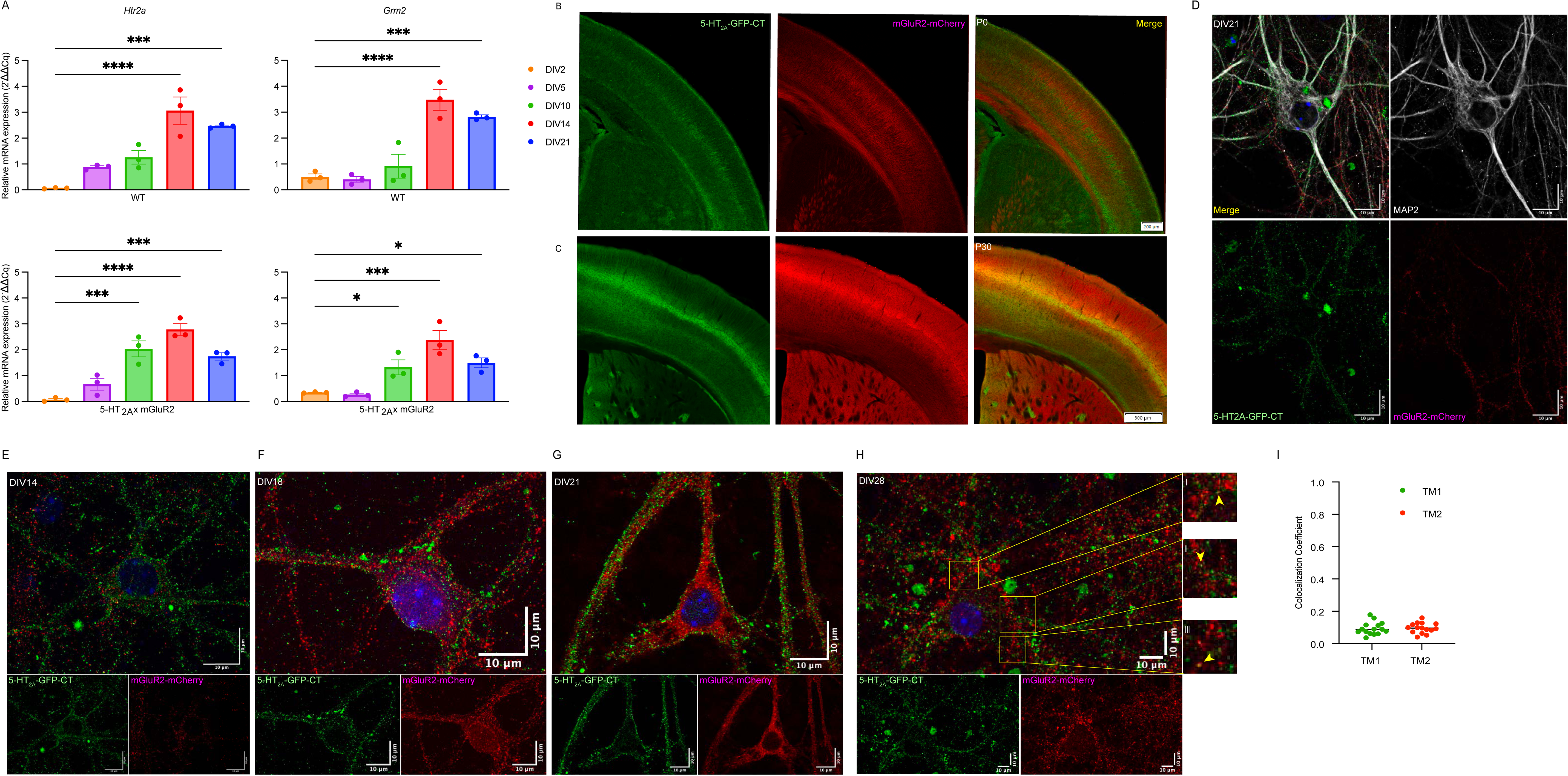
Developmental and subcellular distribution of 5-HT_2A_ and mGluR2 receptors in primary cortical neurons. **(A)** RT-qPCR data for *Htr2a* (left column) and *Grm2* (right column) mRNA expression across DIV 2, 5, 10, 14 and 21 in primary cortical neurons from wild type (WT) C57BL/6J mice (top row) and Htr2a-EGFP-CT x mGluR2-mCherry^+/+^ mice (bottom row). Data represent mean fold-change ± SEM relative to each gene at DIV0. Statistical comparisons by one-way ANOVA with Dunnett’s multiple comparison test from n = 3 independent cultures. \*\*\*\**p < 0.000,* ***p <0.001, **p < 0.01, *p < 0.05, ns = not significant. **(B)** Representative immunofluorescence images of 5-HT_2A_-EGFP-CT (left) and mGluR2-mCherry (middle) expression with merged view (right) from Htr2a-EGFP-CT x mGluR2-mCherry^+/+^ P0 brains slices. Acquired with 10X objective. **(C)** Same as (B) for P30 brain slices. **(D)** Representative DIV21 primary cortical neuron from Htr2a-EGFP-CT x mGluR2-mCherry^+/+^ mice showing co-expression of 5-HT_2A_-EGFP-CT (green), mGluR2-mCherry (red), neuronal marker MAP2 (gray), and DAPI (blue). Images were acquired using a Leica STELLARIS 8 FALCON STED 100X/1.4 NA oil objective and 4X frame averaging with 1X zoom. **(E-G)** Representative images of individual neurons at DIV14 (F), DIV18 (G), and DIV 21(H). Top panels show merged view; bottom panels are 5-HT_2A_-EGFP-CT and mGluR2-mCherry panels, respectively. Images were acquired using a Leica STELLARIS 8 FALCON STED 100X/1.4 NA oil objective and 4X frame averaging with 1X zoom. **(H)** Representative DIV28 neuron. Top panel shows merged view; bottom panels are 5-HT_2A_-EGFP-CT and mGluR2-mCherry panels, respectively. Side panels show zoomed insets corresponding to yellow-boxed regions. Yellow arrows indicate sites of close receptor-receptor proximity in culture. Images were acquired using a Leica STELLARIS 8 FALCON STED 100X/1.4 NA oil objective and 4X frame averaging with 1X zoom. **(I)** Mander’s colocalization coefficient results from double transgenic Htr2a-EGFP-CT × mGluR2-mCherry^+/+^ mice. Colocalization was assessed using the JaCoP plugin in ImageJ and reported as Manders’ overlap coefficients TM1 and TM2, where TM1 represents the fraction of GFP signal overlapping with mCherry signal and TM2 represents the fraction of mCherry signal overlapping with GFP signal. A coefficient of 1 indicates complete overlap and 0 indicates no overlap. Data are expressed as mean ± SEM from 15 images acquired from n = 3 independent cultures (DIV14). Images were acquired using a 100X/1.4 NA oil objective.

To visualize protein expression *in vitro*, primary cortical neurons derived from Htr2a-EGFP-CT x mGluR2-mCherry mice were imaged at DIV14, 18, 21, and 28 using confocal immunofluorescence microscopy. Neuronal identity was confirmed via co-staining with the neuronal marker MAP2, demonstrating that 5-HT_2A_-EGFP positive cells were excitatory neurons (Fig. 6D). Across all timepoints, 5-HT_2A_ receptors were predominantly localized to neuronal plasma membranes, whereas mGluR2 receptors were found in cell bodies and fine neuronal processes (Figs. 6E-H). Thorough colocalization analysis of individual neurons across multiple cultures at DIV14 revealed low Mander’s coefficients (TM1=0.093; TM2=0.096; Fig. 6I), indicating minimal overlap between the GFP and RFP channels.

These results indicate minimal 5-HT_2A_ and mGluR2 receptor colocalization in neurons *in vitro*, although it is possible that 5-HT_2A_ and mGluR2 receptors are in proximity at synapses. To investigate this possibility, high magnification confocal images (Fig. 6H) revealed that mGluR2 and 5-HT_2A_ remain spatially segregated, providing further evidence in favor of the hypothesis that these receptors occupy distinct subcellular compartments. Together, these data indicate that in cultured mouse primary cortical neurons, 5-HT_2A_ and mGluR2 receptors largely occupy separate cellular domains, consistent with minimal physical interaction.

## Discussion

The main finding of this paper is that 5-HT_2A_ and mGluR2 receptors occupy distinct neuronal subdomains supporting a model whereby mGluR2 agonists inhibit the actions of psychedelic 5-HT_2A_ agonists indirectly rather than via heterooligomer formation. Using a variety of orthogonal approaches, we find that 5-HT_2A_ receptors are primarily localized to postsynaptic and somatodendritic compartments of Layer V cortical pyramidal neurons. mGluR2 receptors, by contrast, are found in the neuropil, consistent with a presynaptic localization. Significantly, our studies utilized CRISPR-mediated editing of the endogenous loci to create fluorescently tagged receptors, thereby overcoming limitations of antibody-based approaches, which are often unreliable for GPCRs due to conformational flexibility and high sequence homology. Consistent with previous results, we found mGluR2 and 5-HT_2A_ receptors reach mature expression patterns at similar postnatal timepoints. The relatively late expression of mGluR2 and 5-HT_2A_ receptors is consistent with many prior studies investigating the ontogeny of GPCR expression^72,73^. Using a variety of *in vivo* and *in vitro* approaches, including confocal microscopy, BRET, RAPID nanobody-based purification and proteomics studies, we find no evidence of significant 5-HT_2A_–mGluR2 colocalization or heterodimerization.

In addition to these approaches, our radioligand binding and kinetic analyses provide complementary information regarding potential receptor-level mechanisms. Although prior studies have suggested that activation of mGluR2/3 receptors may alter agonist binding affinity at 5-HT_2A_ receptors^21^, we observed no measurable effect of mGluR2/3 activation on either equilibrium binding parameters or ligand binding kinetics. Furthermore, analysis of mGluR2/3 ligand binding across genotypes demonstrates that the observed biphasic binding profile depends on mGluR2 expression but is independent of 5-HT_2A_ receptor expression. Together, these findings do not support models in which mGluR2 directly modulates 5-HT_2A_ receptor ligand binding or stabilizes distinct affinity states.

Although our RAPID purification failed to detect mGluR2 co-purifying with 5-HT_2A_ receptors, we observed selectively enriched canonical G-protein subunits and synaptic scaffolding proteins, particularly when exposed to the agonist 25CN-NBOH. To maximize the yield of the low-abundance interacting proteins, we used whole-brain homogenates for RAPID isolation, an approach validated by others for enhancing target protein yield^74^. These findings suggest that although there is considerable functional crosstalk between mGluR2 and 5-HT_2A_ receptors^17,75–77^ (Figs. 4H, 4I), physical interactions are not necessary. The absence of detectable effects on ligand binding affinity or kinetics further argues against receptor-level mechanisms involving allosteric modulation of 5-HT_2A_ by mGluR2. Instead, these results are more consistent with models in which mGluR2 signaling influences 5-HT_2A_-dependent responses indirectly through modulation of synaptic glutamate release and circuit-level activity.

Our proteomics studies revealed potential 5-HT_2A_ interactions with protein complexes linked to neuronal plasticity and excitatory neurotransmission, including the AMPA receptor subunit *Gria1*, the voltage-gated sodium channel subunit *Scn4b*, and scaffolding proteins such as *Dock9* and SAP97 (*Dlg1*). Gene ontology analysis highlights roles in membrane potential regulation and action potential processes. While STRING analysis does not confirm direct physical interactions, the observed clustering into discrete biological processes supports context-dependent or indirect signaling assemblies. These results are consistent with prior electrophysiological observations showing that 5-HT_2A_ agonists increase the probability of pyramidal neuronal firing via AMPA receptor mediated EPSCs^78,79^. These findings are also consistent with many prior studies that 5-HT_2A_ agonists enhance excitatory signaling by organizing into protein complexes that facilitate glutamatergic transmission rather than through direct heterodimerization with mGluR2^49–53,60,60,61,63–65^. These observations support hypotheses that psychedelic drugs, via 5-HT_2A_ agonism, induce plasticity^52,80–82^ which has been linked to the putative therapeutic actions of psychedelics^4,83^. Such structural and functional modifications could contribute to the changes in cortical connectivity observed after psychedelic exposure^84,85^.

Functionally, we replicated findings that mGluR2 agonists attenuate the DOI-induced HTR response (Fig. 4I). These findings highlight a clear influence of mGluR2 signaling on 5-HT_2A_ activity, reflecting a decrease in agonist-induced HTR. As comparable effects have been reported for other Gα_i/o_ protein-coupled receptors (e.g. 5-HT_1A_ receptors which do not physically interact with 5-HT_2A_ receptors) which can inhibit both the DOI-induced HTR^86,87^ and electrophysiological responses^19^, functional antagonism can occur in the same neuron without heterodimerization.

Recently several mutually exclusive models of psychedelic drug action have been advanced including: (1) 5-HT_2A_ receptor-mediated activation of Gα_i_ is essential for their psychedelic actions^66^; (2) 5-HT_2A_ receptors are intracellular^88^; (3) 5-HT_2A_ receptors signal via Gα_s_^89^; (4) 5-HT_2A_ receptors signal via β-arrestin^90^; and (5) that 5-HT_2A_ receptors mediate their actions primarily via Gα_q_ pathways^13,46,67,68,91,92^. Our proteomics results do not support models which posit productive 5-HT_2A_ interactions with non-Gα_q/11_ G-proteins as only Gα_q_ and Gα_11_ were enriched while Gα_o_, Gα_i1_ and Gα_z_ were not, and Gα_s_ was undetected under the conditions of our experiments. These proteomic and microscopy studies are also inconsistent with an ‘intracellular’ localization of 5-HT_2A_ receptors in neurons. Instead, our results confirm a large number of prior studies indicating that 5-HT_2A_ receptors are localized to post-synaptic membranes of neurons^25–28,38,51,58^ and the plasma membranes of transfected cells^49,50^ where they couple to Gα_q/11_ G proteins.

Together, these findings support a model in which 5-HT_2A_ and mGluR2 receptors operate independently at the molecular level but converge functionally through compartment-specific interactions. These results have broader implications for dissecting the microcircuit-specific actions of psychedelic compounds and suggest that targeting layer-specific receptor interactions and synaptic protein complexes could provide a novel strategy for the therapeutic modulation of 5-HT_2A_-receptors.

## Acknowledgements

This work was supported by R01MH112205 and R37DA045657 to BLR, and the Brain and Behavior Research Foundation Young Investigator grant (NJW). This research is based in part upon work conducted using the UNC Metabolomics and Proteomics Core Facility, which is supported in part by NCI Center Core Support Grant (2P30CA016086-45) to the UNC Lineberger Comprehensive Cancer Center and Nutrition and Obesity Research Center (P30DK056350). The imaging in this study was conducted using microscopes at the Michael Hooker Imaging Core which is supported by NIH grant 1S10OD030300 (Leica Stellaris 8).

---

**Supplemental Figure 1.**
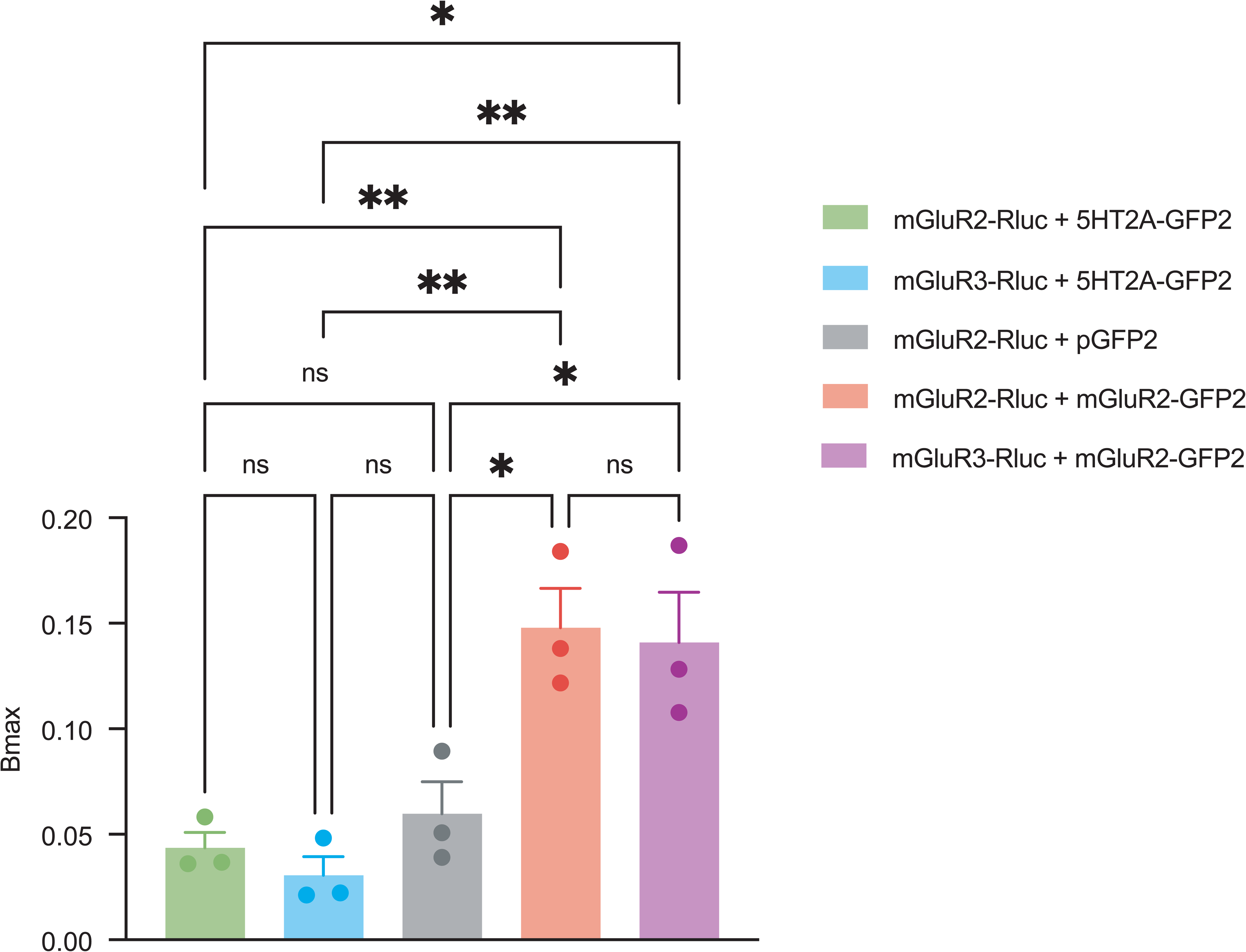
Statistical analysis of receptor-receptor BRET2. Quantification of netBRET2 values from receptor-receptor BRET2 experiments shown in Fig. 1C. netBRET2 values were derived from nonlinear regression curve fitting of BRET2 titration data for each receptor pair (mGluR2-GFP (negative control), mGluR2-mGluR2, mGluR3-mGluR2, 5-HT_2A_-mGluR3, and 5-HT_2A_-mGluR2). The following comparisons were found to be significant: mGluR2-Rluc + 5-HT_2A_-GFP2 vs. mGluR3-Rluc + mGluR2-GFP2, *p = 0.0107; mGluR3-Rluc + 5-HT_2A_-GFP2 vs. mGluR3-Rluc + mGluR2-GFP2, **p = 0.0057; mGluR2-Rluc + 5-HT_2A_-GFP2 vs. mGluR2-Rluc + mGluR2-GFP2, **p = 0.0076; mGluR3-Rluc + 5-HT_2A_-GFP2 vs. mGluR2-Rluc + mGluR2-GFP2, **p = 0.004; mGluR2-Rluc + pGFP2 vs. mGluR3-Rluc + mGluR2-GFP2, *p = 0.0241; mGluR2-Rluc + pGFP2 vs. mGluR2-Rluc + mGluR2-GFP2, *p = 0.0175. Statistical comparisons were performed using one-way ANOVA with Tukey’s multiple comparisons post hoc test. Data are expressed as mean ± SEM from n = 3 independent experiments. *p < 0.05, **p < 0.01.

**Supplemental Figure 2.**
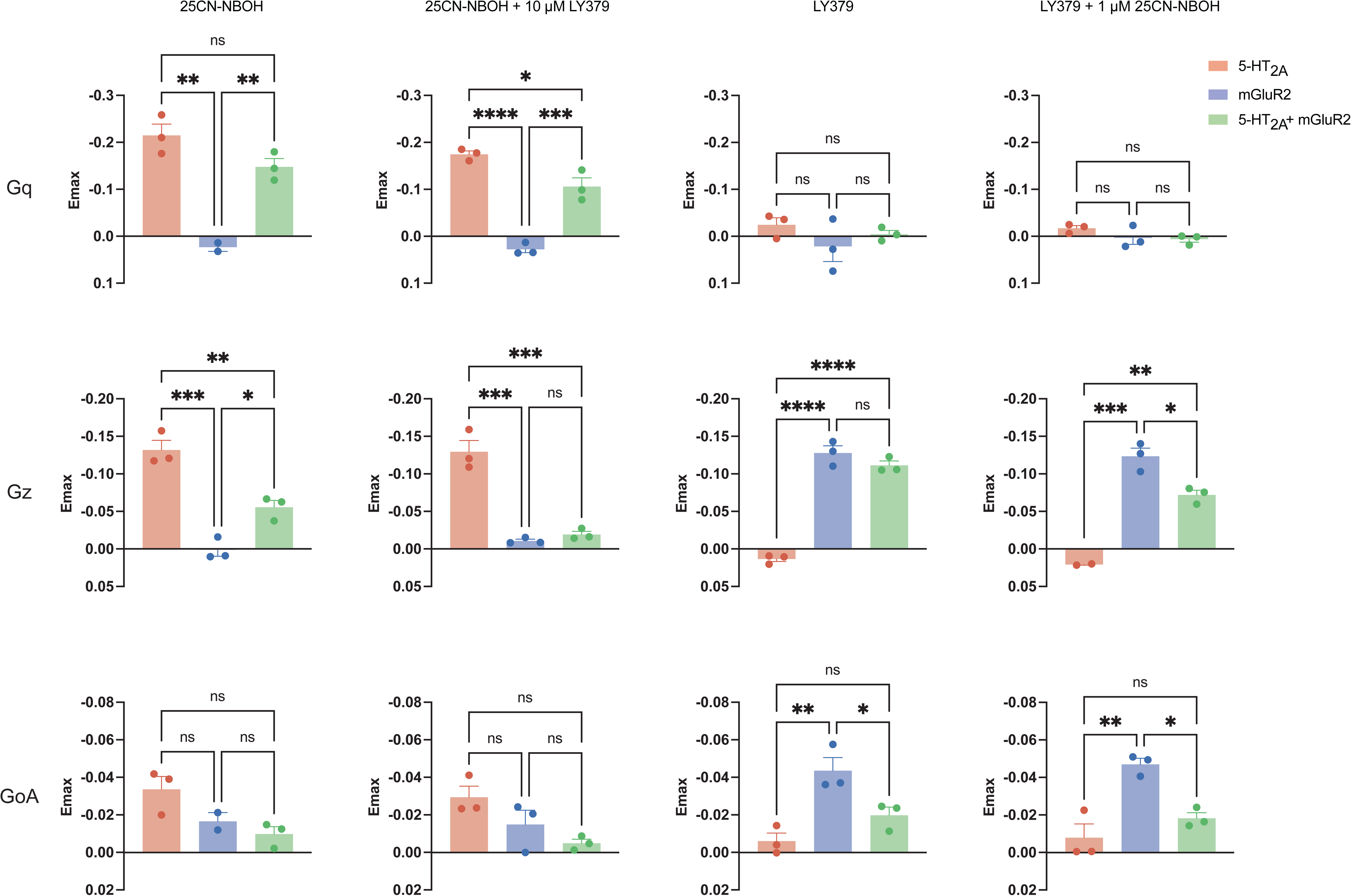
Statistical analysis of G-protein coupling BRET2 signaling assays. Quantification of netBRET2 values from G-protein coupling BRET2 signaling assays shown in Figs. 1E–G. Data are organized by G-protein subunit: Gαq (first row), Gαz (second row), GαoA (third row), and 5-HT_1F_ (fourth row, 5-HT only). For each G-protein, netBRET2 values are shown for cells transfected with 5-HT_2A_ alone (red), mGluR2 alone (blue), or both 5-HT_2A_ and mGluR2 together (green), treated with 25CN-NBOH alone (first column), 25CN-NBOH following 10 μM LY379268 (LY379) pretreatment (second column), LY379 alone (third column), or LY379 following 1 μM 25CN-NBOH pretreatment (fourth column). The following comparisons were found to be significant: Gq/25CN-NBOH – 5-HT_2A_ vs mGluR2, **p = 0.0012; mGluR2 vs 5-HT_2A_+mGluR2, **p = 0.0055; Gq/25CN-NBOH + LY379 – 5-HT_2A_ vs mGluR2, ****p < 0.0001; 5-HT_2A_ vs 5-HT_2A_+mGluR2, ***p = 0.0006; mGluR2 vs 5-HT_2A_+mGluR2, *p = 0.0178; Gz/25CN-NBOH – 5-HT_2A_ vs mGluR2, ***p = 0.0002; 5-HT_2A_ vs 5-HT_2A_+mGluR2, **p = 0.0047; mGluR2 vs 5-HT_2A_+mGluR2, *p = 0.0192; Gz/LY379 – 5-HT_2A_ vs mGluR2, ****p < 0.0001; 5-HT_2A_ vs 5-HT_2A_+mGluR2, ****p < 0.0001; Gz/25CN-NBOH + LY379 – 5-HT_2A_ vs mGluR2, ****p < 0.0001; 5-HT_2A_ vs 5-HT_2A_+mGluR2, ***p = 0.0006; mGluR2 vs 5-HT_2A_+mGluR2, *p = 0.0178; GoA/LY379 – 5-HT_2A_ vs mGluR2, **p = 0.0061; mGluR2 vs 5-HT_2A_+mGluR2, *p = 0.0459; GoA/LY379 + 25CN-NBOH – 5-HT_2A_ vs mGluR2, **p = 0.0033; mGluR2 vs 5-HT_2A_+mGluR2, *p = 0.0146. Statistical comparisons were performed using one-way ANOVA with Tukey’s multiple comparisons post hoc test. Data are expressed as mean ± SEM from n = 3 independent experiments. *p < 0.05, **p < 0.01, ***p <0.001, \*\*\*\**p < 0.0001*.

**Supplemental Figure 3.**
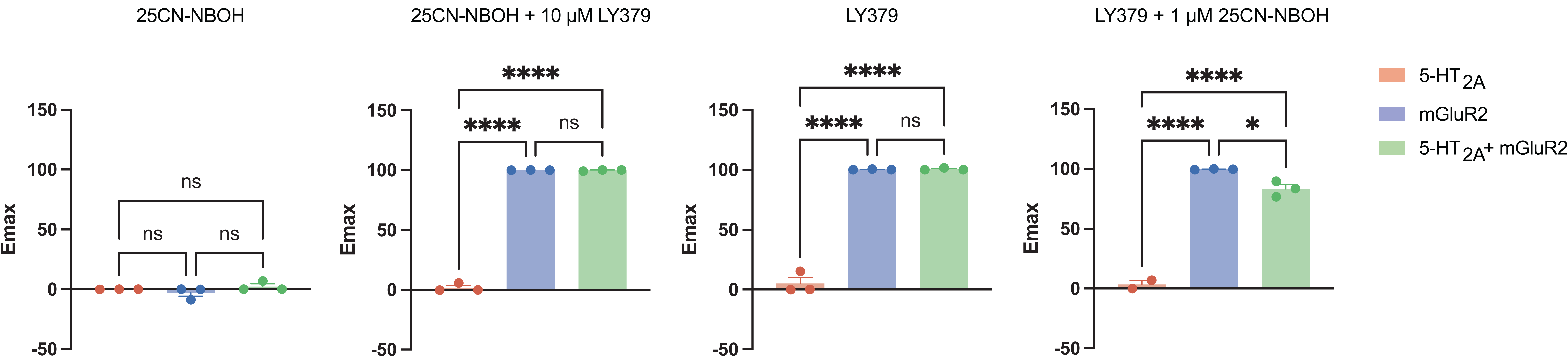
Statistical analysis of GloSensor cAMP assays. Quantification of Imax values from GloSensor cAMP assay data shown in Figs. 1H and 1I. Data are organized by treatment condition from left to right: 25CN-NBOH alone, LY379268 (LY379) alone, 25CN-NBOH following 10 μM LY379 pretreatment, LY379 following 1 μM 25CN-NBOH pretreatment. Imax values (for inhibition conditions) were derived from nonlinear regression using a three-parameter dose-response model. The following comparisons were found to be significant: 25CN-NBOH + LY379 – 5-HT_2A_ vs mGluR2, ****p = 0.5259; 5-HT_2A_ vs 5-HT_2A_+mGluR2, ****p = 0.5097. Statistical comparisons between transfection conditions (5-HT_2A_ alone (red), mGluR2 alone (blue), or both 5-HT_2A_ and mGluR2 together (green)) were performed using one-way ANOVA with Tukey’s multiple comparisons post hoc test. Data are expressed as mean ± SEM from n = 3 independent experiments performed in triplicate. *p < 0.05, **p < 0.01, ***p <0.001, \*\*\*\**p < 0.0001*.

**Supplemental Figure 4.**
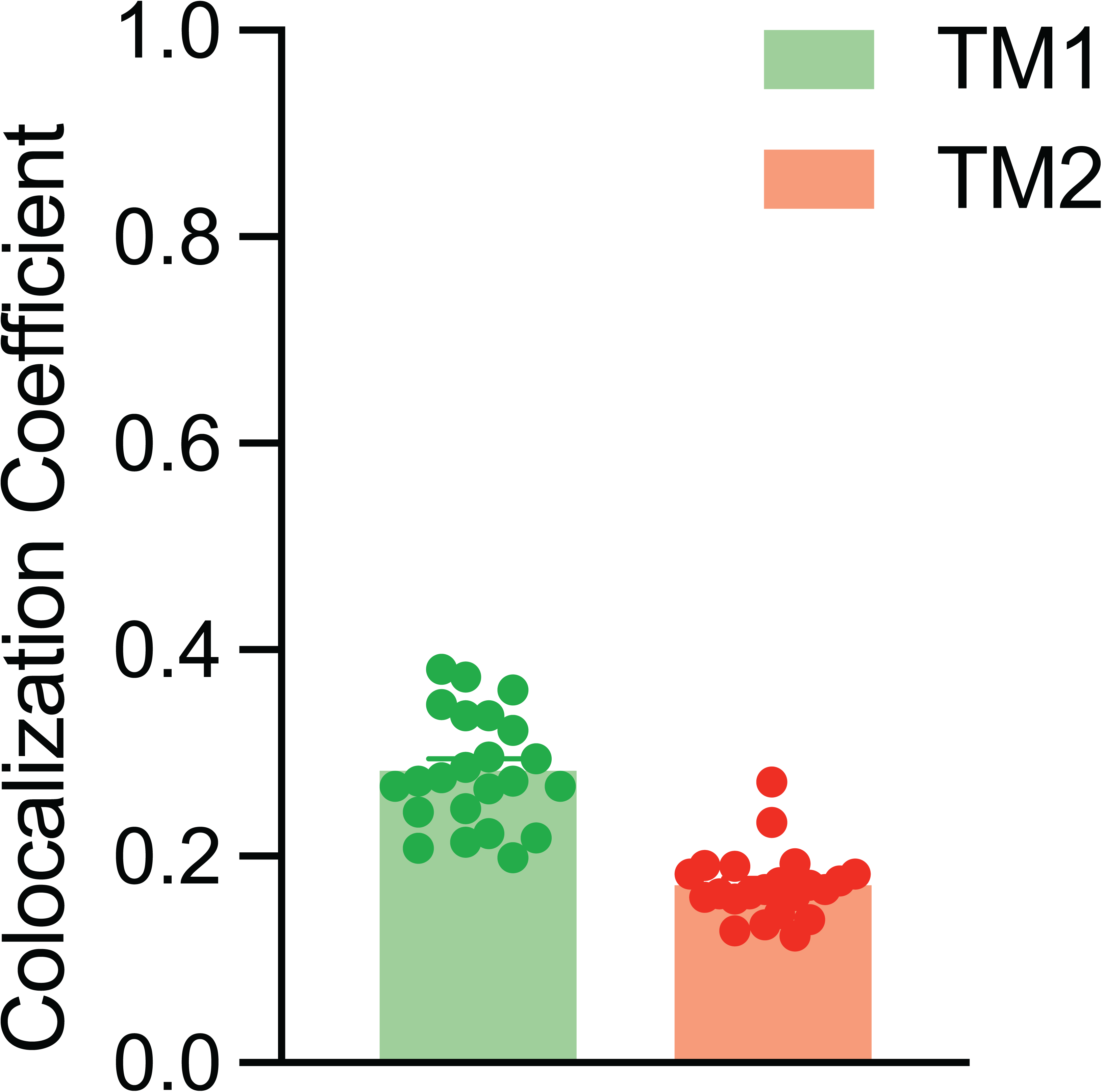
Analysis of in vivo colocalization. Quantification of colocalization between mGluR2-mCherry and 5-HT_2A_-EGFP-CT receptors in brain tissue sections of somatosensory cortex from double transgenic Htr2a-EGFP-CT × mGluR2-mCherry mice. Colocalization was assessed using the JaCoP plugin in ImageJ and reported as Manders’ overlap coefficients TM1 and TM2, where TM1 represents the fraction of GFP signal overlapping with mCherry signal and TM2 represents the fraction of mCherry signal overlapping with GFP signal. A coefficient of 1 indicates complete overlap and 0 indicates no overlap. Data are expressed as mean ± SEM from 7-8 images acquired from each animal (n = 3 animals, P30-P60). Images were acquired using a 100X/1.4 NA oil objective.

